# USP10 strikes down β-catenin by dual-wielding deubiquitinase activity and phase separation potential

**DOI:** 10.1101/2022.10.31.514466

**Authors:** Yinuo Wang, Aihua Mao, Jingwei Liu, Pengjie Li, Shaoqin Zheng, Tong Tong, Zexu Li, Haijiao Zhang, Lanjing Ma, Jiahui Lin, Zhongqiu Pang, Qing Han, Fukang Qi, Xinjun Zhang, Maorong Chen, Xi He, Xi Zhang, Teng Fei, Bi-Feng Liu, Daming Gao, Liu Cao, Qiang Wang, Yiwei Li, Ren Sheng

**Affiliations:** College of Life and Health Science, Northeastern University, Shenyang 110819, China; Division of Cell, Developmental and Integrative Biology, School of Medicine, South China University of Technology, Guangzhou 510006, China; Guangdong Provincial Key Laboratory of Marine Biotechnology, Institute of Marine Sciences, Shantou University, Shantou, Guangdong 515063, China; College of Basic Medical Science, China Medical University, Shenyang 110122, China; Key Laboratory for Biomedical Photonics of MOE at Wuhan National Laboratory for Optoelectronics - Hubei Bioinformatics and Molecular Imaging Key Laboratory, Department of Biomedical Engineering, College of Life Science and Technology, Huazhong University of Science and Technology, Wuhan 430074, China; Key Laboratory of Molecular Biophysics of the Ministry of Education, National Engineering Research Center for Nanomedicine, College of Life Science and Technology, Huazhong University of Science and Technology, Wuhan 430074, China; F.M Kirby Neurobiology Center, Boston Children’s Hospital, Department of Neurology, Harvard Medical School, Boston, MA 02115, USA; College of Sciences, Northeastern University, Shenyang 110004, China; State Key Laboratory of Cell Biology, CAS Center for Excellence in Molecular Cell Science, Shanghai Institute of Biochemistry and Cell Biology, Chinese Academy of Sciences, Shanghai 200031, China

**Keywords:** USP10, ubiquitination, phase separation, β-catenin, colorectal cancer.

## Abstract

Wnt/β-catenin signaling is a conserved pathway crucially governing development, homeostasis and oncogenesis. Discovery of novel regulators holds great values in both basic and translational research. Through screening, we identified a deubiquitinase (DUB) USP10 as a novel and critical modulator of β-catenin. Mechanistically, USP10 binds to key scaffold Axin1 via conserved motifs and stabilizes Axin1 through K48-linked deubiquitination, and surprisingly, tethers Axin1 and β-catenin physically while promoting phase separation for β-catenin suppression regardless of its enzymatic activity. Functionally, USP10 prominently regulates embryonic development and intestinal homeostasis by antagonizing β-catenin via DUB activity. In colorectal cancer, USP10 substantially represses cancer growth mainly through physical binding compensation and phase separation promotion and correlates with Wnt/β-catenin magnitude clinically. Collectively, we discovered USP10 functioning in multiple biological processes against β-catenin and unearthed a novel enzyme-dependent and -independent “dual-regulating” mechanism by which USP10 utilizes parallelly and context-dependently. USP10 inhibitor was suggested in treating certain Wnt-related diseases.

## Introduction

Wnt/β-catenin signaling (aka canonical Wnt signaling) is a pivotally conserved signaling cascade in metazoans.^1, 2^ In vertebrate, Wnt/β-catenin signaling critically governs multiple biological processes including embryonic development and adult tissue homeostasis. Mechanistically, in the absence of Wnt ligand, co-transcription factor β-catenin is contained within the “destruction complex” primarily composed by Axin1 and APC (*Adenomatous Polyposis Coli*), therefore undergoes phosphorylation by GSK3 (Glycogen Synthase Kinase 3)/CK1 (Casein Kinase 1) and ends up with ubiquitination and proteasomal degradation. While Wnt ligand is present, it binds and recruits (co-)receptors to form signalosome on the plasma membrane and disrupts the presence of Axin1 and GSK3 in the destruction complex. As a result, β-catenin becomes released from its confinement, then translocating into the nucleus to initiate the expression of target genes that regulate cell proliferation, differentiation, and self-renewal. Dysregulation of Wnt/β-catenin signaling causes severe developmental disorders, degenerative diseases and malignant tumors, in particular colorectal cancers (CRC).^1, 2^ Though the core of this signaling cascade has been studied for decades, discovery of new regulators is still of great value for both basic and translational medicine. For instance, as novel findings, Rspo and Notum greatly expanded the mechanistic understanding of Wnt signaling regulation, as the antibody or inhibitor against them has been proven promising for particular cancer treatments.^3–9^ As a central component of Wnt/β-catenin signaling, Axin1 is indispensable for both Wnt activation and β-catenin suppression.^1^ It contains RGS domain for APC engagement, a conserved helix for β-catenin binding and DIX domain for multimerization with Dishevelled, while the majority remains unstructured (presumably intrinsically-disordered). However, thanks to this flexibility, Axin1 has been shown capable of either undergoing phase separation or polymerization, which effectively facilitates the entrapment of β-catenin within the destruction complex.^10^ Regulation of Axin1 critically depends on the post-translational modifications (PTM). For instance, the phosphorylation at S497 allows Axin1 to obtain an “open” conformation to make full contact with β-catenin.^11^ Tankyrase (TNKS) can regulate the protein stability by poly-ribosylating Axin1 amino-terminal, which allows Axin1 subsequent ubiquitination by RNF146 for degradation.^12–14^ Despite that the potential toxicity is concerned, TNKS inhibitor such as XAV939, shows promise in treating Wnt-dependent diseases by regulating Axin1 protein level.^12^ Other regulators of Axin1 post-translational modification were also reported, which play important roles in Wnt signaling transduction.^15–19^

Ubiquitination is a protein PTM that rules cellular protein degradation and activities.^20^ By dynamic elongation or removal of K48-linked ubiquitin chain, the fate of protein can be decided for degradation or preservation, which is crucially required for cellular protein homeostasis. Additionally, other residues-linked (K63, K27, and etc.) ubiquitination were known to affect the protein activity. As one of the well-studied deubiquitinases (DUBs), USP10 has been shown to regulate multiple important biological processes.^21^ It was reported that USP10 deubiquitinates substrates including FLT3, PTEN, KLF4 and YAP/TAZ to regulate cancer growth.^22–25^ Other literatures also show that USP10 can deubiquitinate p53, VPS34 complex, NOTCH and AMPK in various biological conditions.^26–29^ The small molecule inhibitor of USP10, Spautin-1, has been shown for translational medical potentials in treating various diseases.^27, 30^

In this study, we have identified USP10 as a critical suppressor of Wnt/β-catenin signaling through cDNA screening. Notably, USP10 dominates Wnt/β-catenin signaling via dual paths. USP10 can directly associate with both β-catenin and Axin1 via conserved motifs and remarkably stabilize Axin1 through K48-specific deubiquitination. And unexpectedly, USP10 can also corroborate the puncta formation of Axin1 through physical constraining-mediated phase separation, which is independent of its DUB activity. In terms of biological functions, we find that USP10 plays crucial roles in embryonic development and intestinal homeostasis by modulating the amplitude of Wnt/β-catenin signaling via DUB activity. And in human CRC, USP10 significantly represses tumor progression mainly through enzymatic-independent function both *in vitro* and *in vivo*, and correlates with patient survival and Wnt magnitude clinically. Thus, the Axin1-stabilizing and phase separating paths work parallelly and context-dependently. Taken together, we have identified that USP10 directly regulates Wnt/β-catenin signaling for the first time and revealed a novel DUB-dependent and -independent “dual-wielding” mechanism. By employing multiple biological models, we have elucidated the functional significance of USP10 on Wnt regulation, and suggest the therapeutic promise for treating developmental and regenerative defects by using USP10 inhibitor.

## Results

### USP10 is a negative regulator of Wnt/β-catenin signaling

We performed DUBs cDNA screening under Wnt3a conditioned medium (CM) stimulation by exercising TOPFlash reporter (B/R system) in HEK293T cells as readout to identify novel DUBs that potentially regulate Wnt/β-catenin signaling. To obtain reliable results, we used two different transfection levels of cDNA as two independent experiment sets and included stringent controls in the screen. Amongst the initially screened 30 DUBs, we found that USP10 had a strong inhibitory effect on Wnt/β-catenin signaling which was comparable to that of Axin1 (**Fig.1A; S1A**). This inhibitory effect of ectopically expressed USP10 on Wnt/β-catenin signaling was then validated in human CRC cells such as RKO and DLD-1 (**Fig.S1B-C**). To further consolidate the results, we showed that USP10 inhibited Wnt/β-catenin signaling in a dose-dependent manner presented by both the luciferase assay and cytosolic β-catenin level (**Fig.1B-C**). Knockout or knockdown of USP10, on the contrary, activated Wnt/β-catenin signaling as shown in various assays, including TOPFlash reporter and the accumulation of cytoplasm β-catenin (**Fig.1D-E; S1D-H**). To exclude the possibility that USP10 affects Wnt/β-catenin through other related pathways, we employed dual-luciferase chemiluminescence to detect the changes of target genes including TGFβ, BMP, CREB and etc. The result showed that expression of USP10 had little effect on these pathways, thus strongly suggested that USP10 exerts suppression on Wnt/β-catenin directly (**Fig.S1I**).

**Figure 1.**
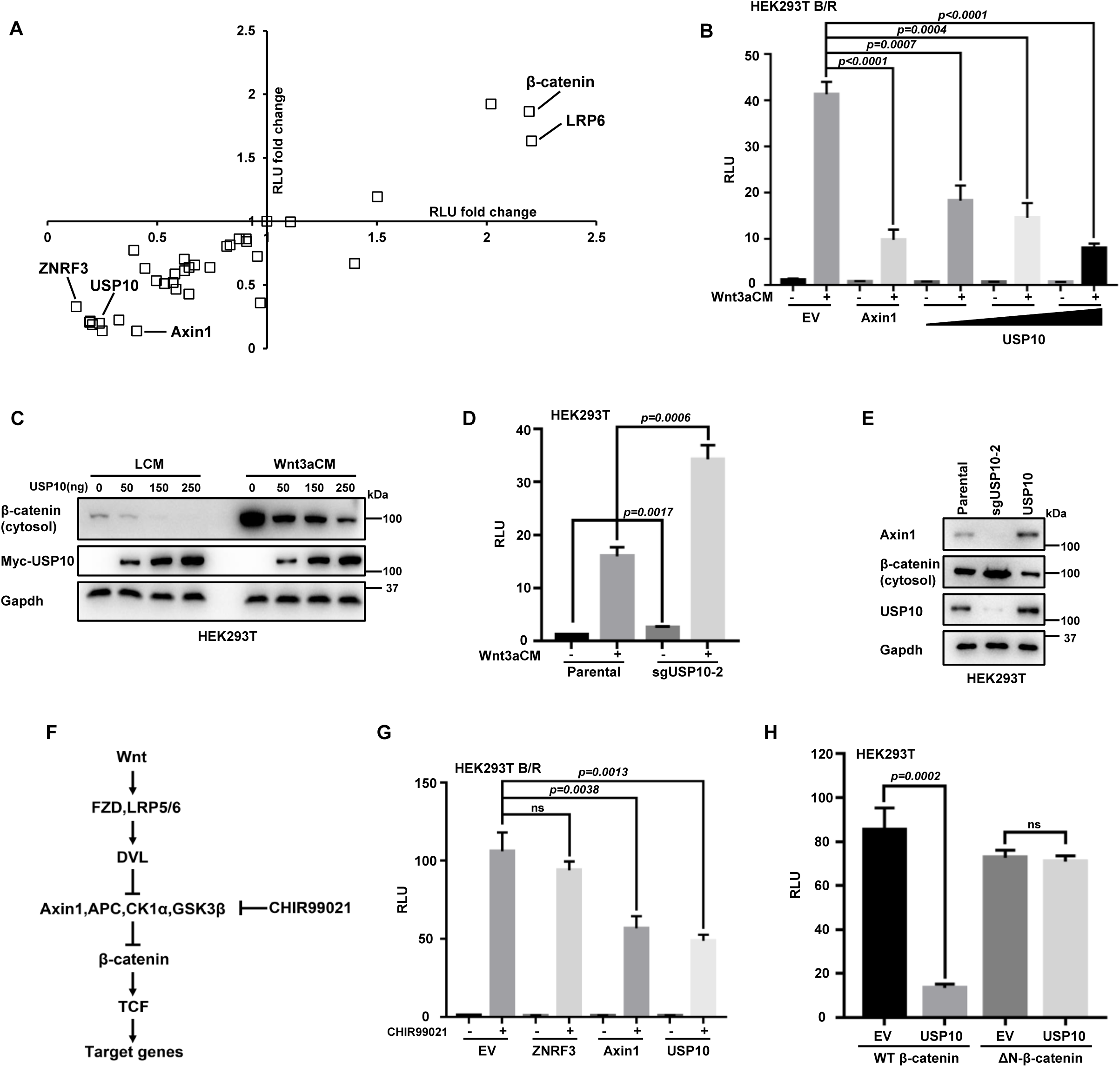
USP10 is a negative regulator of Wnt/β-catenin signaling. (A) DUB cDNA screening on Wnt/β-catenin signaling by TOPFlash reporter fold changes in HEK293T cell. The two axes represent two independent sets of experiments. LRP6 and β-catenin are positive controls, while ZNRF3 and Axin1 are negative controls. (B) USP10 significantly reduces TOPFlash activity fold change dose-dependently at the presence of Wnt3a CM. Error bars mean ± SD, n = 3, two-tailed Student’s t-test. (C) USP10 reduces cytosolic β-catenin accumulation dose-dependently shown by WB assay. (D) Knockout of USP10 significantly enhances TOPFlash reporter at the presence of Wnt3a CM. Error bars mean ± SD, n = 3, two-tailed Student’s t-test. (E) WB assay showing the alteration of cytosolic β-catenin levels under USP10 overexpression or knockout condition. (F) Schematic flow of Wnt/β-catenin signal transduction process. (G) TOPFlash reporter assay under the treatment of GSK3 inhibitor CHIR99021. ZNRF3: Wnt signalosome control. Axin1: β-catenin destruction complex control. Error bars mean ± SD, n = 3, two-tailed Student’s t-test. (H) TOPFlash reporter assay under the expression of wild-type and ΔN-β-catenin. Error bars mean ± SD, n = 3, two-tailed Student’s t-test. RLU: Relative Luciferase Unit. ns, not significant.

Next, we sought to elucidate the mechanism. To effectively unravel the complexity of Wnt/β-catenin signaling, we first used GSK3 inhibitor CHIR99021 to divide Wnt/β-catenin cascade into the upstream (Wnt signalosome formation) and the downstream events^31–33^ (β-catenin destruction complex formation and afterwards) (**Fig.1F**). After treatment of CHIR99021, USP10 still showed significant inhibition on Wnt/β-catenin signaling that was comparable to Axin1, indicating that the manner of USP10 functioning was independent of Wnt signalosome formation (**Fig.1G**). Meanwhile, we constructed the N-terminal truncated β-catenin (ΔN-β-catenin), a mutant commonly used as the constitutively active variant due to absence of GSK3 and CK1 phosphorylation sites.^34^ By comparing the wild-type (WT) β-catenin and ΔN-β-catenin, we showed that USP10 was only able to effectively inhibit Wnt/β-catenin signaling during WT β-catenin overexpression but not ΔN-β-catenin, which indicated that USP10 could not significantly affect β-catenin nuclear transportation (**Fig.1H; S1J**). Taken together, we have identified USP10 as a novel negative regulator of Wnt/β-catenin signaling and argued that USP10 functions at the β-catenin destruction complex level.

### USP10 stabilizes Axin1 through K48-linked deubiquitination

Next, we examined the interaction of USP10 with the key components of the destruction complex (including β-catenin, Axin1, APC, GSK3 and β-Trcp). By employing co-immunoprecipitation (co-IP), we found that USP10 co-existed with β-catenin, Axin1 and GSK3 in the same complex at both exogenous and endogenous levels (**Fig.2A; S2A**). Since USP10 was a repressor of Wnt/β-catenin signaling, we intuitively focused on Axin1 and logically hypothesized that USP10 might regulate Axin1 ubiquitination and stability. We first showed that endogenous Axin1 protein level could be upregulated upon treatment of the proteasome inhibitor MG132 (**Fig.2B**). At the presence of ectopic USP10, the stability of Axin1 was enhanced regardless of MG132 while Axin1 transcription level remained unaltered (**Fig.2B; S2B**). Interestingly, the presence of USP10 showed even higher Axin1 protein level than protease inhibitor, which suggested USP10 might have other functions beyond Axin1 deubiquitination. This Axin1-increasing effect was also dose-dependent on USP10 level (**Fig.S2C**). To check the protein stability of Axin1, pulse-chase assay was exercised by blocking the *de novo* protein synthesis and observing the turnover of the existing protein. We found that USP10 overexpression enhanced endogenous Axin1 stability, as USP10 depletion showed the opposite effect (**Fig.2C; S2D-E**). The inhibitory effect of USP10 on Wnt/β-catenin was strongly attenuated when Axin1 was knocked-down, indicating that USP10 regulates Wnt/β-catenin signaling mainly through Axin1 (**Fig.2D; S2F**).

**Figure 2.**
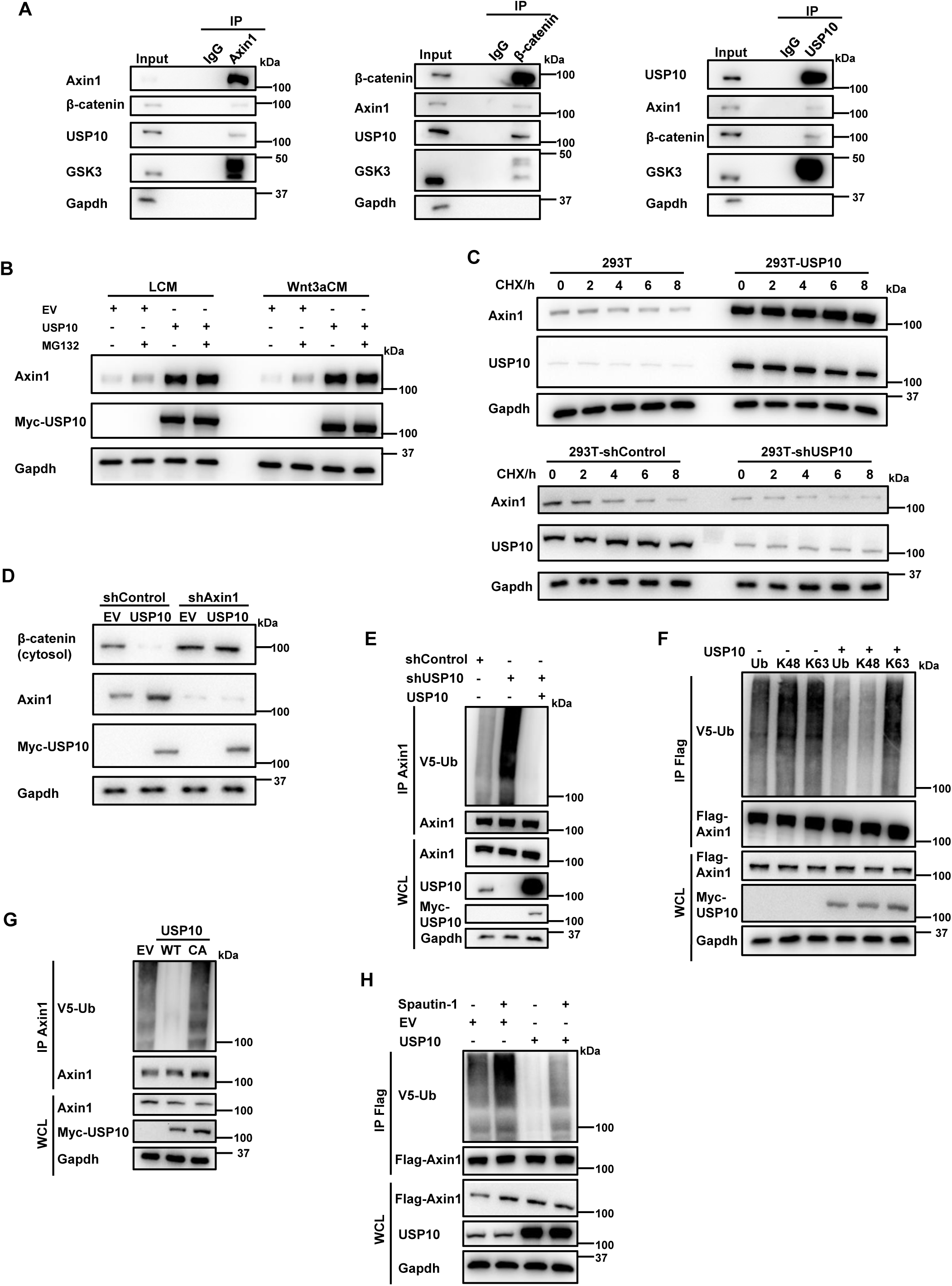
USP10 stabilizes Axin1 through K48-linked deubiquitination. (A) Co-IP assay showing USP10 interaction with Axin1, β-catenin and GSK3 endogenously. (B) Axin1 protein level changes under the expression of USP10 or/and the treatment of proteasome inhibitor MG132. (C) Pulse-chase assay of endogenous Axin1 under USP10 overexpression (upper) or depletion (lower) conditions. (D) Axin1 and cytosolic β-catenin levels under Axin1 knockdown or/and USP10 overexpression condition. (E) Ubiquitination assay of Axin1 under USP10 knockdown and rescue conditions. (F) Ubiquitination assay showing USP10 mainly affects Axin1 K48-ubiquitination. (G) Ubiquitination assay showing USP10-CA mutant loses the capability to deubiquitinate endogenous Axin1. (H) Ubiquitination assay showing USP10 inhibitor Spautin-1 effectively thwarts USP10 DUB activity and increases Axin1 ubiquitination level.

In terms of Axin1 ubiquitination, USP10 depletion resulted in aggravated Axin1 ubiquitination, which could be recovered by USP10 rescue (**Fig.2E; S2G**). It was further confirmed that Axin1 was a direct substrate of USP10 via *in vitro* deubiquitination assay (**Fig.S2H**). Moreover, USP10 mainly removed K48-linked ubiquitination on Axin1, which was consistent with its regulation on Axin1 stability (**Fig.2F**). The effect was lost when we employed USP10-CA (USP10 C424A, enzymatically-dead mutant)^27^, suggesting USP10 deubiquitinates Axin1 based on its DUB activity (**Fig.2G; S2H-I**). Based on the binding results, we also showed USP10 could not deubiquitinate β-catenin (**Fig.S2J**). This result brought clear evidence that USP10 serves unambiguously as Wnt/β-catenin antagonist, unlike another DUB, USP7, which was contradictorily reported to stabilize both Axin1 and β-catenin.^15, 35^ On the other hand, we employed the USP10 inhibitor Spautin-1 to validate our results through chemical perturbation. The addition of Spautin-1 enhanced the ubiquitination of Axin1, destabilized Axin1 and inhibited the activity of the Wnt/β-catenin, which drew consistent conclusions with the genetic manipulations (**Fig.2H; S2K, S2L**).

### USP10 acts as a scaffold in the destruction complex by connecting Axin1 and β-catenin

We then mapped the binding site between USP10 and Axin1. Topologically, USP10 contains two major parts: a conserved DUB domain at its carboxyl terminal as the rest of the protein forms one lengthy non-structured region at its amino terminal (named as ΔDUB) (**Fig.3A**). We therefore constructed and expressed USP10 full-length (FL), CA, DUB and ΔDUB *in vitro* and performed pull-down experiments. It was clearly seen that USP10 FL, CA and ΔDUB interacted with Axin1 as USP10 DUB domain alone lost the binding capacity (**Fig.3B; S3A**). Since the ΔDUB was unstructured, we arbitrarily divided it into four pieces with approximate 100 amino acids for each, named Segment 1-4 (S1-S4), respectively (**Fig.3A**). By employing the pull-down and co-IP assays, we found that S2 was mainly responsible for Axin1 binding (**Fig.3C; S3B**). Since Wnt/β-catenin signaling is highly conserved, we argued that the binding between USP10 and Axin1 should involve in their conserved residues. Through the alignment of USP10 in different species, we found seven conserved motifs within the ΔDUB (**Fig.S3C**). Through co-IP and pull-down assays, we confirmed the USP10 binding site was a conserved polybasic region (PBR) in S2 region (a.a. 143-163) (**Fig.3D; S3D**).

**Figure 3.**
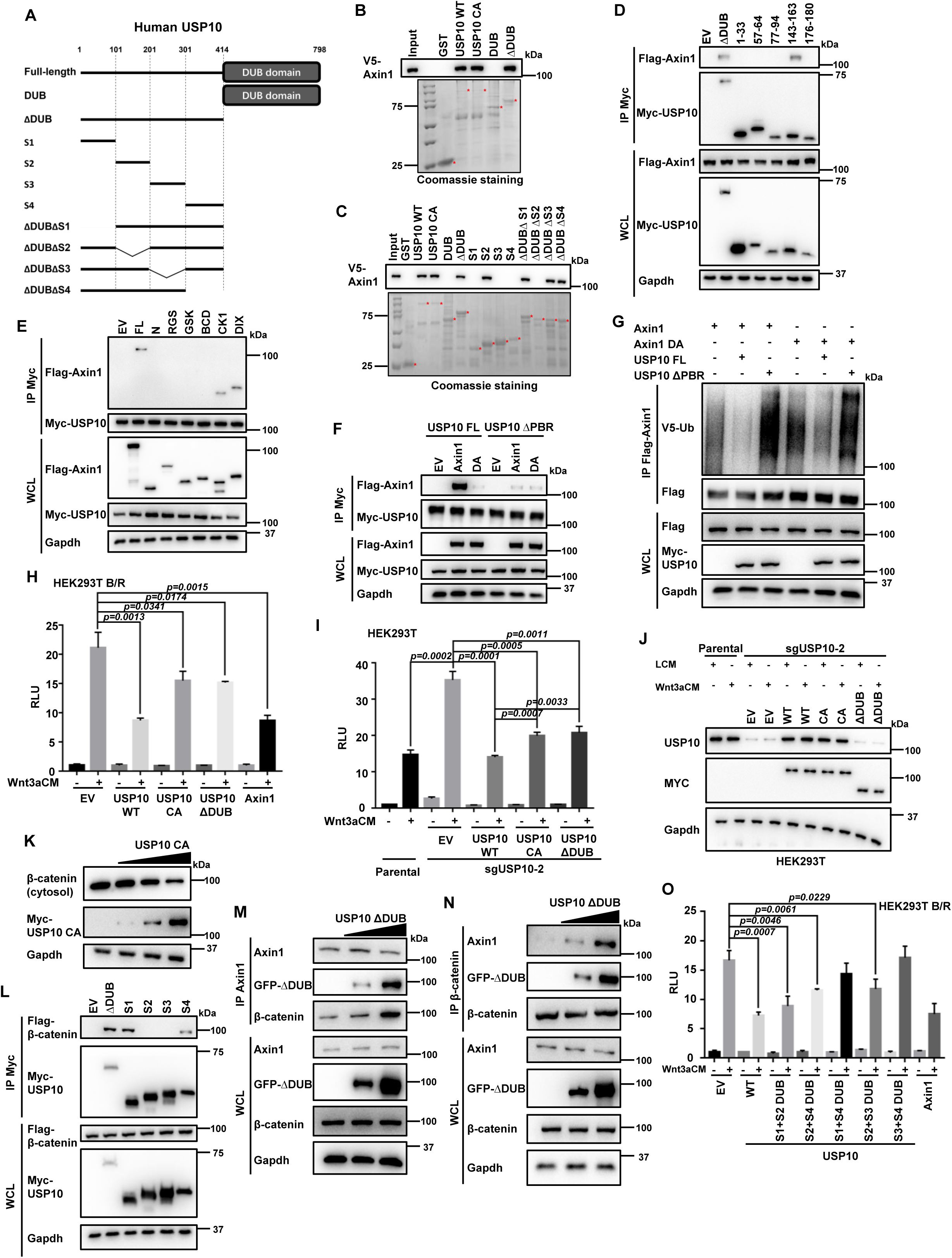
USP10 acts as a scaffold in the destruction complex by connecting Axin1 and β-catenin. (A) Schematic diagram of USP10 truncation mutants constructed in this work. (B, C) *In vitro* pull-down assay of different GST-USP10 fragments with Axin1. Asterisks represent the major bands of the desired proteins. (D) Co-IP assay of Axin1 with different conserved regions of USP10. (E) Co-IP assay of USP10 with different segments of Axin1. (F) Co-IP assay of USP10 WT/ΔPBR with Axin1 WT/DA. (G) Ubiquitination assay showing USP10-ΔPBR cannot deubiquitinate Axin1. And Axin1-DA cannot be deubiquitinated by USP10. (H, I) TOPFlash reporter assay showing USP10-CA and USP10-ΔDUB retain moderate inhibitory effect on Wnt/β-catenin by overexpression (H) or USP10 knockout and rescue (I). Error bars mean ± SD, n = 3, two-tailed Student’s t-test. (J) USP10 protein level test by WB in (I). The endogenous USP10 antibody cannot identify ΔDUB region and thus immunoblotting on Myc serves as a ruler for protein level. (K) WB showing cytosolic β-catenin level diminishes dose-dependently of USP10-CA. (L) Co-IP assay showing different segments of USP10 interacting with β-catenin. (M, N) Co-IP assay showing the interaction strength of endogenous Axin1 and β-catenin enhances while the dose of USP10-ΔDUB increases. (O) TOPFlash reporter assay showing the minimal functional truncations of USP10 require S2 region with the DUB domain. Error bars mean ± SD, n = 3, two-tailed Student’s t-test.

We further divided Axin1 into six parts based on earlier study (**Fig.S3E**).^11^ From co-IP and pull-down assays, we found that USP10 mainly interacted with the putative CK1 binding region and DIX domain (**Fig.3E; S3F**). It was previously report that Axin1 DIX domain contains a negatively-charged patch that forms self-inhibition intramolecularly.^11^ Thus, we hypothesized that this patch was responsible for USP10 PBR binding through electrostatic interaction. We observed that the binding between Axin1 and USP10 was significantly alleviated when we mutated the negatively-charged patch to neutral residues (Axin1-DA) or we truncated the USP10 PBR (ΔPBR) (**Fig.3F**). It was further confirmed that losing either the PBR on USP10 or the negatively-charged patch on Axin1 caused loss-of-function effect of USP10 in Axin1 ubiquitination and stabilization (**Fig.3G; S3G**). Taken together, USP10 directly binds to Axin1 through its conserved PBR. Axin1 is a direct substrate of USP10 for K48-linked deubiquitination which leads to the ensuing Axin1 stabilization.

While elucidating the deubiquitination of Axin1 by USP10, we noticed another interesting phenomenon. Despite the lack of DUB activity, expression of USP10-CA and USP10-ΔDUB consistently reduced TOPFlash reporter activity in a moderate manner instead of behaving dominant-negative (**Fig.3H**). And from the knockout and rescue experiment, the same conclusion was drawn (**Fig.3I-J**). We thus hypothesized USP10 might affected Wnt/β-catenin signaling besides Axin1 deubiquitination. First, we verified that neither USP10-CA nor -ΔDUB could deubiquitinate Axin1 (**Fig.S3H**). However, USP10-CA and -ΔDUB could reduce cytoplasmic β-catenin in a dose-dependent manner (**Fig.3K; S3I**). Together with the finding that USP10 interacted with multiple components of the β-catenin destruction complex, we proposed that USP10-ΔDUB may function as a scaffold that physically enhances the complex formation like Amer1/WTX.^36–38^ When we looked into the USP10-β-catenin interaction, we found USP10 S1 and S4 were responsible, which primarily bound to 1-5 and 9-12 Armadillo repeats on β-catenin (**Fig.3L; S3J-M**). Gradient transfection of ΔDUB led to increased interaction between Axin1 and β-catenin dose-dependently (**Fig.3M-N**). USP10 depletion, on the other hand, reduced the binding between Axin1 and β-catenin endogenously (**Fig.S3N**). In brief, we proposed that USP10 bound to Axin1 mainly by the conserved residues in S2 and to β-catenin by S1 and S4. S2 is absolutely required for both DUB-dependent and -independent activity of USP10 as it is the only region capable of Axin1 binding. S1 and S4 are responsible for β-catenin engagement for DUB-independent activity of USP10, which are not required for Axin1 deubiquitination. In line with our hypothesis, the minimal inhibitory units and dominant negative mutants were validated by the luciferase assays (**Fig.3O; S3O**).

### USP10 facilitates the puncta formation of Axin1 via a IDRs-mediated phase separation-like manner independent of DUB activity

After we dissected the contribution of USP10 in the physical interaction between Axin1 and β-catenin, we investigated the stabilization of Axin1 granule. The formation of Axin1 granule was supposed to form via either IDR (intrinsically disordered region) - mediated liquid-liquid phase separation or DIX domain-mediated oligomerization in previous literatures.^10, 39, 40^ As we tested above, the interaction between the Axin1 and USP10 happened between the DIX domain and PBR of USP10 via multivalent electrostatic interactions. Briefly, the PBR of USP10 locates within a lengthy non-structured IDR, while electrically neutralized DIX domain efficiently prevented the interaction without affecting the original function of the DIX domain^11^; these results suggested that the stabilization of USP10 on Axin1 granule taken place mechanistically through an IDRs-mediated phase separation-like process rather than DIX domain mediated heat-to-tail polymerization. Meanwhile, we want to note that the physical phase separation and polymerization/oligomerization favors each other^41–43^, which might augment the stabilization of Axin1 granule in the downstream. Nevertheless, our live imaging of Axin1-mCherry clearly showed their dynamic fusion and splitting of the formed droplet-like structure (**Fig.S4A**). A fluorescence recovery after photobleaching (FRAP) assay further confirmed the granules of Axin1 maintained its recovery capability (**Fig.S4B-C**).^44^ From the assay of FRAP, we showed that the Axin1 granule contained both the recoverable fraction and immobile fraction, which suggested it contained both the DIX domain-mediated polymerization and phase separation from physical interaction.

To study whether USP10 could regulate the granule formation of Axin1, we simultaneously expressed USP10 and Axin1 (**Fig.4A**). Co-expression with USP10 led to more Axin1 granules per cell together with larger sizes as compared to Axin1 transfection alone (**Fig.4B-C**). In addition, we analyzed the immobile faction of Axin1 in the granules using FRAP assay to reflect the irreversibility in the Axin1-mCherry granules. A larger immobile fraction indicated that stabler Axin1 granules were obtained with overexpressed USP10 (**Fig.4D**). An opposite trend was obtained when we depleted the endogenous USP10 by shRNA (**Fig.4B-D**). To further confirm that it was the non-structured region-mediated phase separation assisting the granule formation of Axin1, we showed that USP10-ΔDUB still promoted the granule formation of Axin1, which confirmed that the importance of the physical interaction in its function. As noted, unlike the USP10 FL, USP10-ΔDUB was not able to deubiquitinate and elevate the cellular contents of the Axin1, thus, USP10-ΔDUB elevated the granule formation of Axin1 fully on its non-structured region-mediated phase separation instead of increasing the concentration of Axin1 at the first place. More significantly, USP10-ΔDUB exhibited an even stronger promotion effect on the granule formation of Axin1 compared to the full-length USP10 as indicated by the diameter (**Fig.4C**), which might suggest the non-structured region-mediated phase separation has a more direct effect as compared to the deubiquitinating functionality. This result clearly indicated that the ΔDUB was sufficient for the granule augmentation via IDR-mediated phase separation. Consistently, USP10-CA stabilized the granules of Axin1 as shown by their larger size (**Fig.4C**) and their increased immobile fraction under FRAP (**Fig.4D**), despite of the decreased numbers of Axin1 granule per cells formed as compared to USP10 WT (**Fig.4B**). One could argue that USP10-CA was more efficient to coalesce small Axin1 puncta into larger ones.

**Figure 4.**
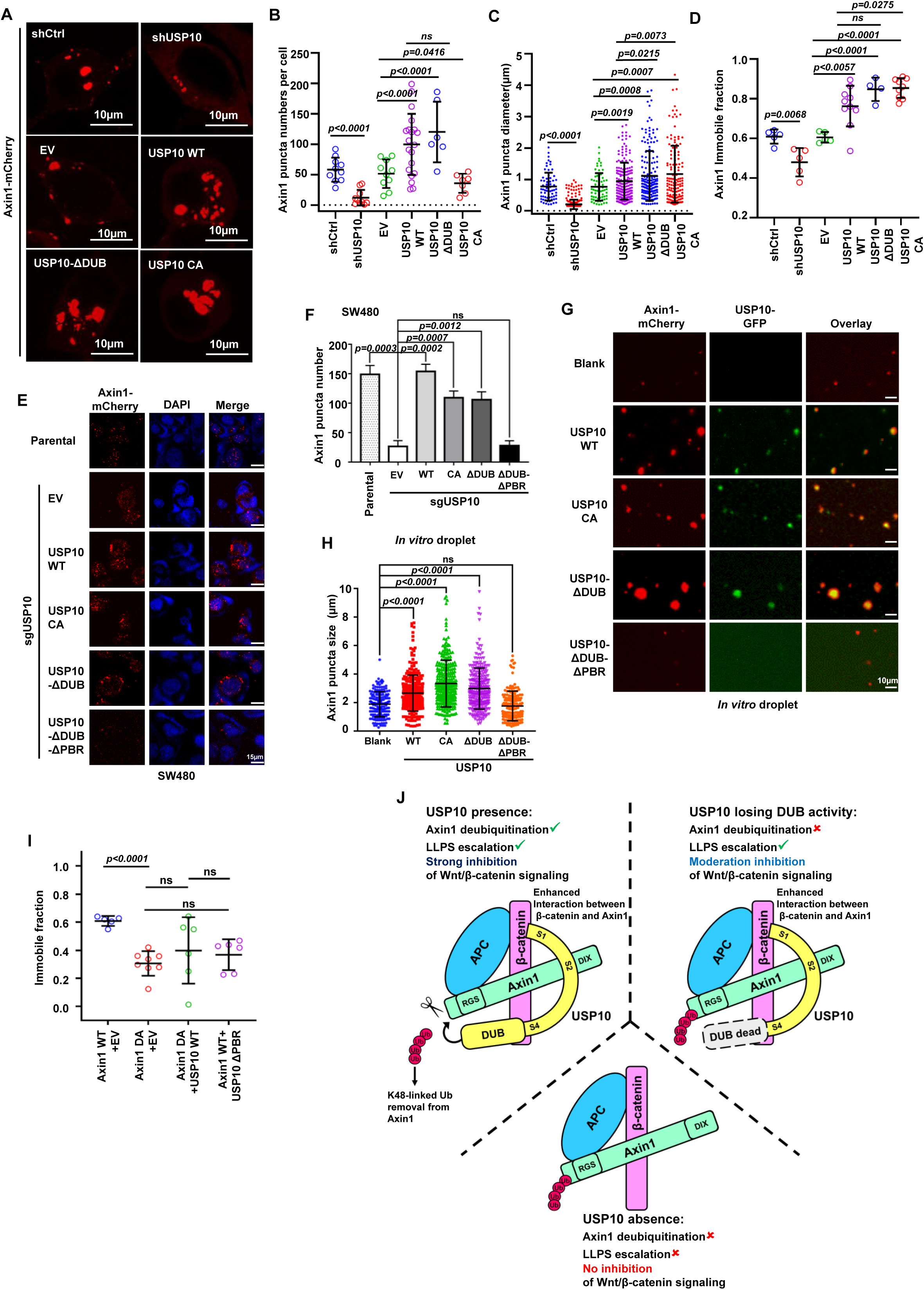
USP10 facilitates the puncta formation of Axin1 via a IDRs-mediated phase separation-like manner independent of DUB activity. (A) Representative fluorescent images of Axin1 droplets when co-expressed with shCtrl, shUSP10, EV, USP10 WT, USP10 CA and USP10-ΔDUB. (B-D) Numbers of Axin1 puncta per cell (B), size of Axin1 puncta (C), and immobile fraction (D) of Axin1 puncta. Error bars mean ± SD, two-tailed Student’s t-test. (E) Representative figures of Axin1 puncta by immunostaining of endogenous Axin1 in SW480 cell. Red, Axin1(Alexa 555). Blue, DAPI. All figures are in the same scale in this panel. (F) The statistical analysis of (E). Error bars mean ± SD, by two-tailed Student’s t-test. (G) Representative figures of *in vitro* phase separation assay by co-incubation of bacterial expressed Axin1-mCherry and USP10-GFP (WT and mutants). Red, Axin1-mCherry. Green, USP10-GFP. USP10 ΔDUB ΔPBR does not colocalize with Axin1 and thus appears as widespread background in green channel. All figures are in the same scale in this panel. (H) The statistical analysis of (G). Error bars mean ± SD, by two-tailed Student’s t-test. (I) Immobile fraction of Axin1-DA puncta when co-expressed with USP10 WT and USP10-ΔPBR, as compared to Axin1 WT puncta. (J) Working model of USP10 inhibiting Wnt/β-catenin signaling. The DUB activity contributes to deubiquitination and stabilization of Axin1, and the unstructured region promotes LLPS of the destruction complex through physical interactions. RLU: Relative Luciferase Unit. ns, not significant.

We also exercised additional assays to substantiate the findings. First, we employed immunofluorescence to observe the endogenous puncta formation of Axin1. The experiment was performed in SW480 cells, in which endogenous Axin1 formed puncta as described by the previous literature.^45^ We knocked out the endogenous USP10 by CRISPR-Cas9 and rescued with Myc-tagged WT or mutant USP10 to the approximately equal level of the endogenous protein (**Fig.S4D-E**). The results illustrated that in the absence of USP10, Axin1 puncta number was significantly reduced (**Fig.4E-F**). Rescue by either USP10 WT, CA or ΔDUB could recover the puncta formation of endogenous Axin1 (**Fig.4E-F**). Also, we performed *in vitro* droplet formation assay to further dissect the role of USP10 in regulating Axin1 phase separation. We expressed mCherry-tagged Axin1 and GFP-tagged USP10 (WT and mutants) in *E. coli* and purified the proteins by chromatography. It was clearly seen that Axin1 droplet size significantly enlarged after co-incubation with USP10 WT, CA or ΔDUB (**Fig.4G-H**). The fluorescent microscopy also showed strong co-existence of the red and green channels. As the negative control, ΔDUB ΔPBR (internal truncation of PBR in ΔDUB region) neither could coexist with Axin1 nor promote Axin1 droplet formation (**Fig.4E-H**). Together, we conclude that the USP10 stabilizes the puncta of Axin1 via phase separation independent of its DUB activity but relying on the intrinsically-disordered region.

Next, we tested whether the enhancement effect of USP10 on Axin1 granule via phase separation was through the multivalent physical interactions. As we mentioned above, Axin1-DA, which contains a negative-charged patch in DIX domain turning into neutral residues, exhibited a significantly suppressed binding to USP10 (**Fig.3G**). As compared to Axin1 WT, we further showed that the number, the diameter and the stability of Axin1-DA puncta were significantly lessened (**Fig.S4F-H; Fig.4I**). More importantly, combination of either USP10 WT with Axin1-DA or USP10-ΔPBR with Axin1 WT no longer promotes the formation of puncta, as no phase separation between Axin1 and USP10 can be elevated after eliminating the multivalent physical interactions (**Fig.4I**). Thus, our results suggested that USP10-Axin1 physical interactions are mainly responsible for the enhancement of granule via phase separation. Based on these finding, we argue that USP10 directly augments the Axin1-containing puncta through its IDRs-mediated phase separation, which could enhance the formation of β-catenin destruction complex in the downstream regardless of the DUB activity.^10^ We proposed a mechanistic model that USP10 may repress Wnt/β-catenin signaling via two parallel paths. USP10 can stabilize Axin1 through its DUB activity and meanwhile physically connect the key components in the β-catenin destruction complex through a phase separation-like effect (**Fig.4J**). Each path individually or synergistically contributes to the repressive effect of USP10 on Wnt/β-catenin signaling.

### USP10 functions in embryonic dorsoventral patterning and axis formation through Wnt/β-catenin signaling

The potential roles of USP10 in embryonic development are poorly understood. Zebrafish has emerged as a powerful model to investigate embryonic development due to its external fertilization and transparent embryos. To gain insight into the functions of USP10 during zebrafish embryonic development, we firstly examined its expression by whole-mount *in situ* hybridization with an antisense probe. As shown in (**Fig.S5A**), zebrafish *usp10* transcript was ubiquitously detected in embryos from the one-cell stage up to the early somite stage. Interestingly, the expression of *usp10* was predominantly found in the developing eye field at 24 hours post-fertilization (hpf), which was remarkable decreased at 36 hpf. These results indicate that *usp10* is a maternal and zygotic gene and might have potential roles in early embryonic development. We then assessed the effects of human USP10 (hUSP10) overexpression on embryogenesis. Over 80% of the embryos injected with wild-type hUSP10 mRNA displayed ventralized phenotypes at 24 hpf, including a variably reduced head and notochord and expanded ventral tissues (**Fig.5A-B**).

**Figure 5.**
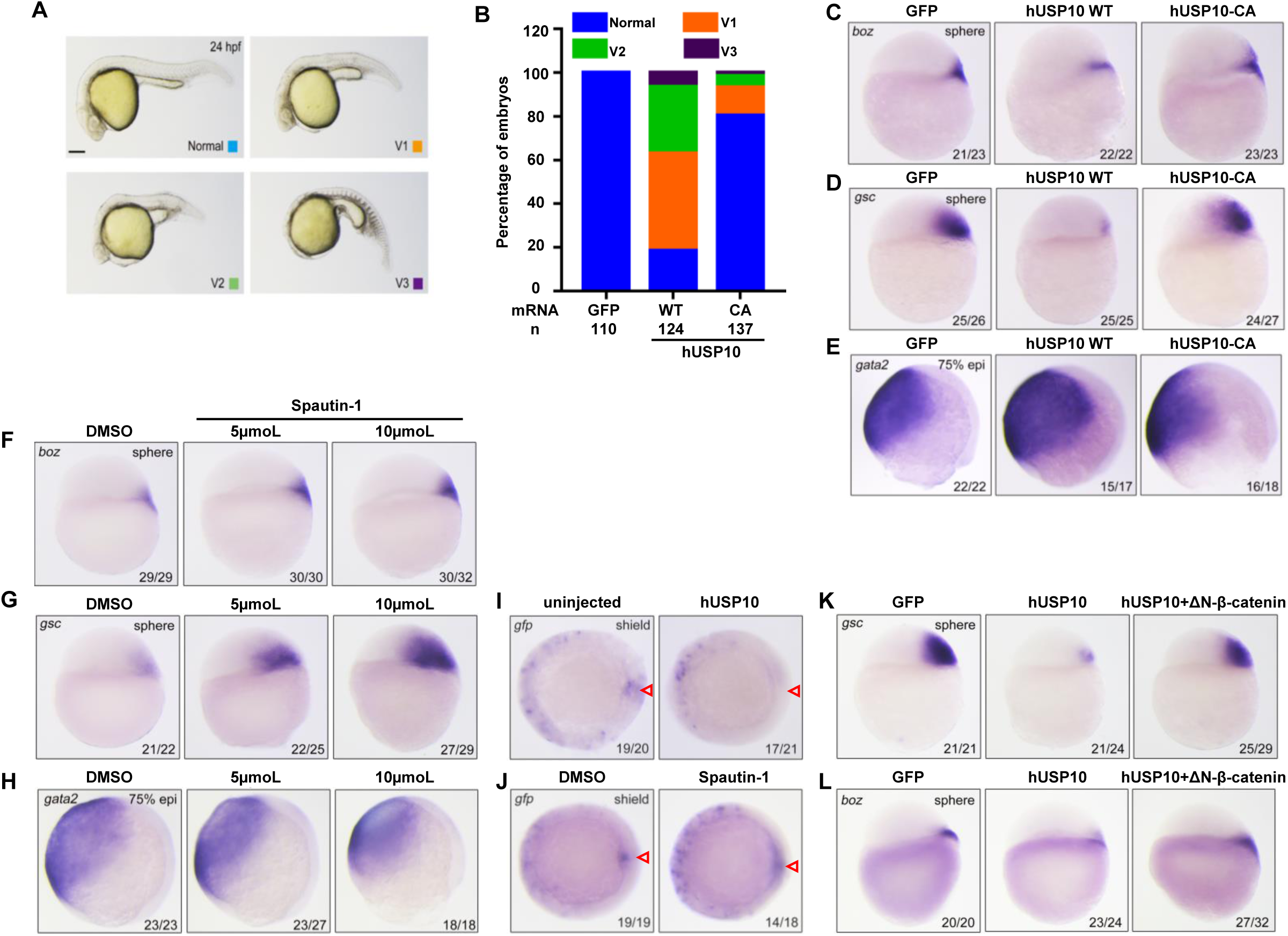
USP10 functions in embryonic dorsoventral patterning and axis formation through Wnt/β-catenin signaling. (A, B) Embryos were injected with 500pg hUSP10 or hUSP10-CA mRNA at the one-cell stage. Representative embryos of different classes at 24 hpf were shown in (A), lateral views with anterior to the left. The percentage of embryos with indicated phenotype were shown in (B). Scale bar, 200μm. (C-E) The expression analysis of dorsal marker genes (C, *boz* and D, *gsc*) at the sphere stage and ventral marker gene (E, *gata2*) in embryos injected with indicated mRNA at the 75% epi (epiboly) stage. Lateral views. Embryos were injected with 500pg hUSP10 or hUSP10-CA mRNA at one-cell stage, 500pg GFP injection was used as a control. (F-H) Dorsal marker genes (F, *boz* and G, *gsc*) and ventral marker gene (H, *gata2*) were assessed in DMSO and Spautin-1 (5μmoL or 10μmoL) incubated embryos at indicated stage by *in situ* hybridization. Lateral views. (I, J) Whole-mount *in situ* hybridization analyzed the transcript of gfp in hUSP10-injected (I) or Spautin-1 (J) treated *Tg(TOPdGFP)* embryos at shield stage. Animal views with dorsal side to the right. Red arrows point to the dorsal organizer of the embryo. (K, L) Overexpression of *ΔN-β-catenin* mRNA rescued hUSP10-induced ventralization. Embryos were injected with 500pg of hUSP10 mRNA alone or together with 100pg of *ΔN-β-catenin* mRNA at the one-cell stage and harvested at the sphere stage for *in situ* hybridization with the probe of *gsc* (K) and *boz* (L). The number of the embryos was indicated within each panel.

To further confirm the role of USP10 in the embryonic dorsal-ventral pattering, the expression of dorsal markers, such as *bozozok (boz)* and *goosecoid* (*gsc*) were analyzed at the sphere stage by *in situ* hybridization. We found that overexpression of hUSP10 led to decreased expression of *boz* and *gsc* (**Fig.5C-D**). Meanwhile, the expansion of ventral non-neural ectoderm (indicated by *gata2* expression) further revealed the dorsal-ventral defects in hUSP10 overexpressed embryos (**Fig.5E**). On the contrary, expression of the DUB-dead mutant, USP10-CA, showed minor effect as compared to WT USP10 (**Fig.5B-E**). Inhibition of the DUB activity of endogenous USP10 in wild-type embryos by treatments with Spautin-1 obviously strengthened the expression of dorsal marker genes and repressed the expression of ventral gene *gata2* in a dose-dependent manner (**Fig.5F-H**). These results suggest that USP10 functions in embryonic dorsal-ventral patterning mainly via its DUB activity.

Previous studies have demonstrated that maternal Wnt/β-catenin signaling is essential for dorsal-ventral patterning during embryogenesis.^46–48^ Given that our findings from cell culture systems indicated that USP10 inhibits Wnt signaling, we asked whether USP10 would affect embryonic dorsoventral axis formation through regulating Wnt/β-catenin pathway. To do this, hUSP10 mRNA was injected into transgenic embryos expressing GFP reporter under TCF/LEF/β-catenin responsive promoter (TOPdGFP).^49^ Results from *in situ* hybridization experiments revealed that hUSP10 overexpression remarkably reduced the GFP reporter expression in the dorsal organizer, suggesting a defective activation of Wnt/β-catenin signaling (**Fig.5I**). On the other hand, blocking the DUB activities of endogenous USP10 in *Tg(TOPdGFP)* embryos by Spautin-1 treatment strongly enhanced GFP expression (**Fig.5J**).

Furthermore, the reduced expression of dorsal marker genes *gsc* and *boz* in hUSP10 overexpressed embryos were well restored by co-injection with *ΔN*-*β*-*catenin* mRNA (**Fig.5K-L**). These data suggested that USP10 has a profound impact on maternal Wnt/β-catenin signaling during early embryonic development. In addition, consistent with the data *in vitro*, immunoblotting results showed an obvious increase of endogenous zebrafish Axin1 protein level in embryos with hUSP10 overexpression (**Fig.S5B-C**). In contrast, inhibition of the deubiquitinate activity of USP10 resulted in a significant decrease of Axin1 (**Fig.S5D-E**). Altogether, these results indicate that USP10 has a conserved role in regulating Axin1 stability depending on its DUB activity, which is responsible for proper dorsoventral axis patterning during embryo development.

### USP10 critically regulates intestinal organoid homeostasis

Intestinal organoid (mini-gut) culturing emerges as an essential tool to study intestinal homeostasis *ex vivo.*^50–52^ It is a three-dimensional structure that consists intestinal-specific cell types and cell lineage and can recapitulate the growth and differentiation of intestinal epithelium. The sphere-like organoid represents the clustering of Lgr5^+^ intestinal stem cells and the budded organoid represents the well-differentiated epithelial mini-structure. It is well established that intestinal organoid growth and differentiation are highly dependent on Wnt/β-catenin signaling amplitude.^50, 52^ In intestinal “stem cell” state culture (sphere-like), we initially found that the organoid could not survive from the overexpression of USP10, which prevented us from statistical analysis. We therefore employed the “Tet-off” system to study the effect.

When doxycycline was removed from the medium, we found that ectopic expression of USP10 led to the straight intestinal stem cell death similar to the effect of Axin1, and this phenotype could be rescued by the addition of GSK3 inhibitor (**Fig.6A-B**). Under such circumstance, overexpression of USP10-CA only showed partial effect in resulting cell death (**Fig.6A-B**). When USP10 was depleted in differentiated intestinal organoid culture (budded), the organoid architecture changed as the number of sphere-like bodies was markedly increased (**Fig.6C-D; S6A-B**). This change of stemness was further evidenced by the increase of proliferation marker Ki67 and reduction of differentiation marker Krt20 (**Fig.6C**). And this effect could be effectively reversed by the addition of TNKS inhibitor XAV939, which accumulates Axin1 to counteract USP10-loss (**Fig.6C-D)**. While we used Spautin-1 to inhibit USP10 DUB activity in intestinal organoid culture, we noticed a significant increase in the number of sphere-like bodies which represented the increase of self-renewal potential (**Fig.6E-F; S6C-D**). However, the diameters of Spautin-1 treated spheres were significantly smaller than those of GSK3 inhibitor, which indicated their difference in potentiating stem cell growth and suggested the scaffolding and phase-transitional function of USP10 could have potential influence in this situation (**Fig.6G; S6E-I**). Together, these results indicate that USP10 modulates intestinal organoid growth and differentiation *ex vivo* through Wnt/β-catenin signaling. Both the DUB activity and biophysical property of USP10 contribute to the intestinal organoid homeostasis. USP10 inhibitor such as Spautin-1 possesses therapeutic potential in regeneration for Wnt/β-catenin stimulation.

**Figure 6.**
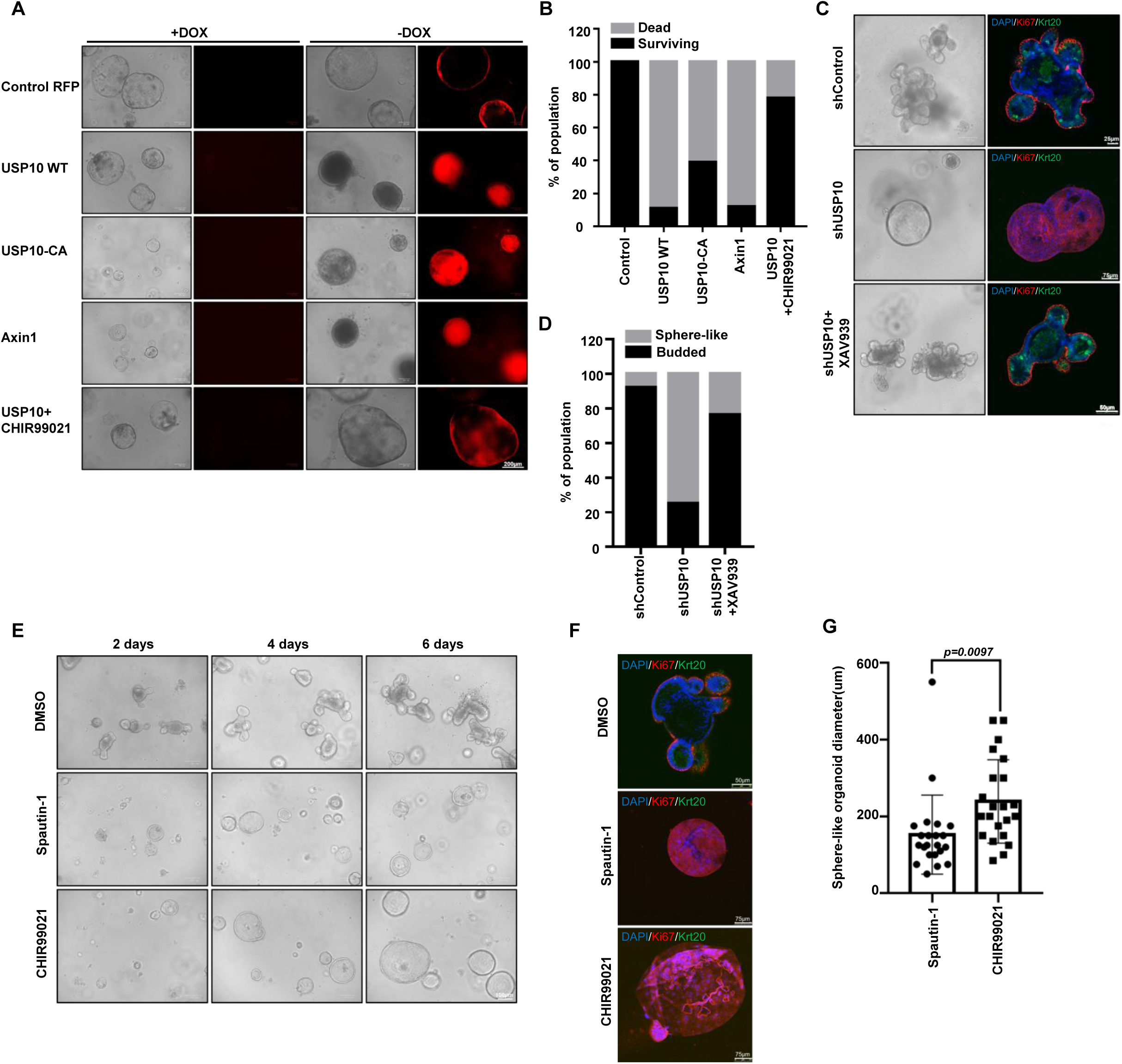
USP10 critically regulates intestinal organoid homeostasis. (A) Representative images (brightfield and red fluorescence) of murine intestinal organoids under different conditions. Dox: doxycycline. (B) Quantitative analysis of the surviving/dead organoids percentage in a. Control group:n=65, USP10 WT group:n=71, USP10 CA group:n=67, Axin1 group:n=67, USP10 WT+CHIR99021 group:n=60. (C) Representative images (brightfield and immunofluorescence) of murine intestinal organoids under USP10 depletion and XAV939 addition. Blue, DAPI; red, Ki67; green, Krt20. (D) Quantitative analysis of the surviving/dead organoids percentage in c. shControl group:n=66, shUSP10 group:n=67, shUSP10+XAV939 group:n=68. (E, F) Representative images (brightfield and immunofluorescence) of murine intestinal organoids under DMSO, Spautin-1 or CHIR99021 treatment. Blue, DAPI; red, Ki67; green, Krt20. (G) Organoid (sphere-like) diameters quantifications in E shown by histogram. Error bars mean ± SD, n = 22, by two-tailed Student’s t-test. All images in any individual panel are in the same amplification scale except the individually marked ones.

### USP10 suppresses CRC growth through inhibition of Wnt/β-catenin signaling

Wnt/β-catenin dysregulation, in particular APC-truncation is the predominant cause for human CRC.^53^ We thus assessed the role of USP10 in CRC. We first verified that in APC-truncated CRC cell lines, ectopic expression of USP10 effectively increased endogenous Axin1 level and reduced cytosolic β-catenin amount, as depletion of USP10 behaved oppositely (**Fig.7A-C; S7A**). Functionally, we found that ectopic expression of USP10 significantly inhibited CRC cell growth both two- and three-dimensionally, as well as cancer cell migration, whereas USP10 depletion (both knockout and knockdown) or overexpression of the dominant-negative USP10 mutants accelerated cancer growth instead (**Fig.7D-G; S7B-F**). Notably, expression of the DUB-dead mutant, USP10-CA, was capable to effectively mitigate the cancer growth comparable to the effect of USP10 WT (**Fig.7D, 7F-G**). Molecularly, USP10-CA reduced the cytosolic β-catenin level and Wnt-target genes (Wnt feedback: AXIN2 and LGR5; cell proliferation: MYC and CCND1) to the same extent of USP10 WT (**Fig. 7H-I**). This data demonstrates the enhancement effect of USP10 on physical phase separation but not USP10 DUB activity is mainly required to withhold CRC growth. Considering that either loss of Axin1 or APC can cause the release of β-catenin ^1^, we reason that in APC-truncated CRC the compensation of APC scaffolding function by USP10 is predominantly required, whereas the stabilization of Axin1 plays a lessor role (**Fig.7J**). And we also logically argue that the USP10-mediated anti-tumor effect in CRC may largely depend on its inhibition on Wnt/β-catenin instead of other pathways, given the fact that all other tumor-suppressing USP10 functions known previously rely on its DUB activity.^21, 54^

**Figure 7.**
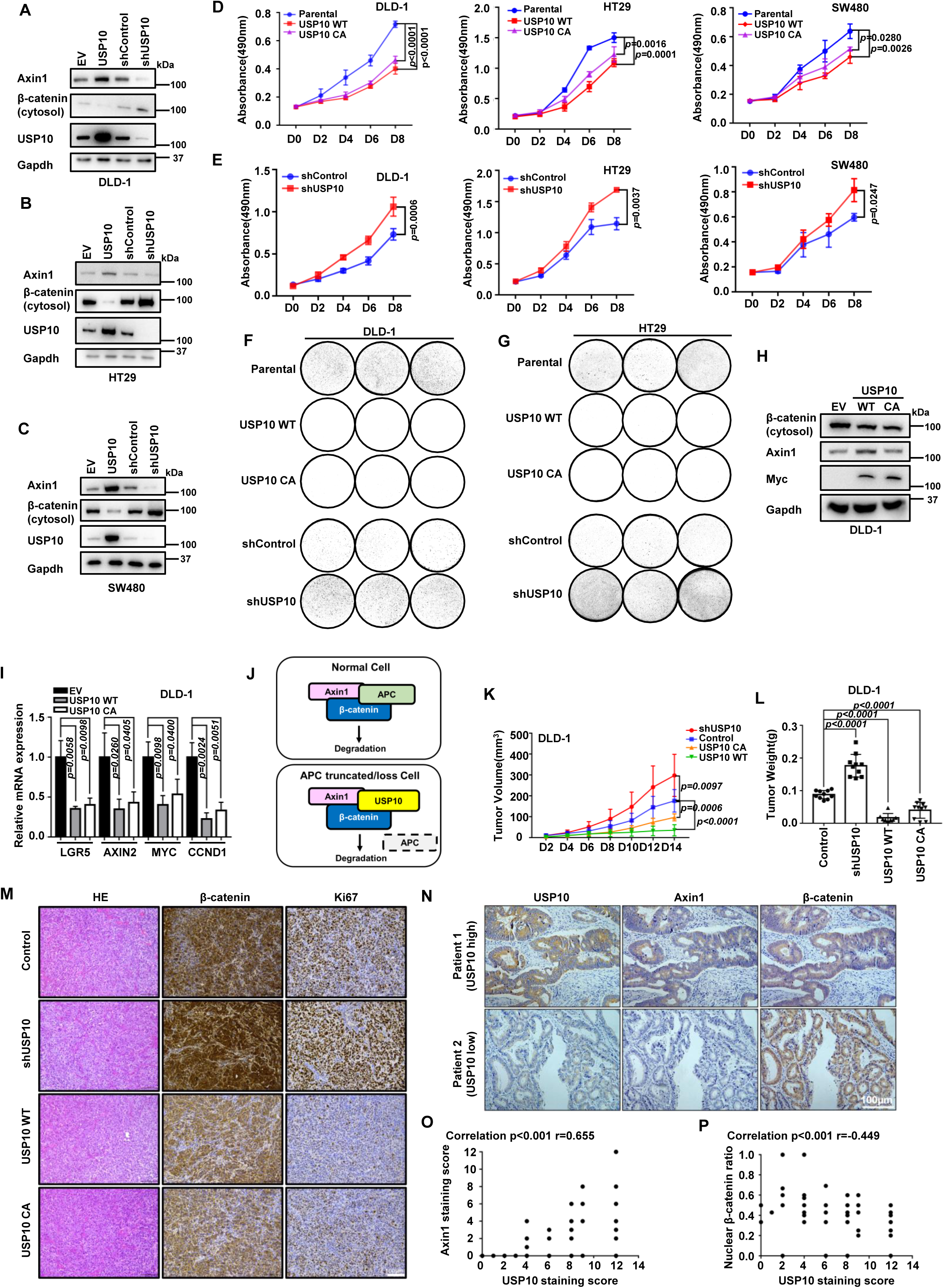
USP10 suppresses CRC growth through inhibition of Wnt/β-catenin signaling. (A-C) Endogenous Axin1 and cytosolic β-catenin levels in CRC cell lines under USP10 overexpression or depletion condition. (A) DLD-1, (B) HT29, (C) SW480. (D, E) MTT cell growth assay of DLD-1, HT29 and SW480 cells under USP10 WT and USP10-CA overexpression (D) or USP10 depletion condition (E). Error bars mean ± SD, n = 3, two-way ANOVA. (F, G) Colony formation assays of DLD-1 (F) and HT29 (G) under USP10 WT/CA overexpression or USP10 depletion condition. (H) Endogenous Axin1 and cytosolic β-catenin levels in DLD-1 cells with overexpression of WT USP10 and USP10-CA. (I) RT-qPCR assay of Wnt target genes expression, including LGR5, AXIN2, MYC and CCND1 in DLD-1. Error bars mean ± SD, n = 3, two-tailed Student’s t-test. (J) Proposed model of USP10 function in cells with loss-of-function APC. USP10 predominantly compensate the scaffolding effect of APC by physically binding to both Axin1 and β-catenin and promoting phase separation. (K, L) Quantitative analyses of the tumor volume (K) and weight (L) of subcutaneously transplanted DLD-1 cells. Tumor volume: Error bars mean ± SD, n = 10 for each group, by two-way ANOVA analysis. Tumor weight: Error bars mean ± SD, n = 10 for each group, by two-tailed Student’s t-test analysis. (M) Representative IHC staining images of β-catenin and Ki67 of the tumor formed from subcutaneously transplanted DLD-1 cell. (N) Representative IHC staining images of Axin1 and β-catenin in CRC patients in USP10-high (upper row) and USP10-low (lower row) groups. (O, P) The correlations of USP10 with Axin1 (O) and nuclear localized β-catenin ratio (P) in CRC patients. By two-tailed Spearman correlation analysis. All images in any individual panel are in the same amplification scale. ns, not significant.

We next transplanted these cells subcutaneously onto nude mice for *in vivo* study. For the transplanted DLD-1 cells, the size and weight of the derived tumors were significantly increased when USP10 was depleted (**Fig.7K-L; S7G**). In contrast, either USP10 WT or USP10-CA overexpression caused significant diminish of the *in vivo* tumor growth, which was consistent with the *in vitro* results (**Fig.7K-L; S7G**). The immunohistochemistry (IHC) showed stronger staining for Ki67 and β-catenin in shUSP10 group than the control group (**Fig.7M**). And the intensities of these staining were attenuated when either USP10 WT or USP10-CA was ectopically expressed (**Fig. 7M**).

To further consolidate the role of USP10 in CRC, we looked into the clinical database and acquired the human tumor specimens. From TCGA database, we found that USP10 expression showed significant positive correlation with longer overall survival in CRC patients (**Fig.S7H**). This correlation in prognosis was also supported by another research which studied a Korean CRC cohort.^55^ To determine the consequence of differential expression of USP10 in human CRC, 92 primary CRC tissues on a microarray were examined by IHC. Representative images were shown in **Fig.7N**. The grouped analysis based on USP10 intensity showed clear difference in Axin1 and nuclear β-catenin ratio in the CRC samples (**Fig.S7I-J**). Overall, higher USP10 level significantly correlated with higher Axin1 level and lower nuclear-localized β-catenin ratio, which was fully supportive to our molecular and cell biological studies (**Fig.7N-P**). Collectively, our data has shown that USP10 suppresses CRC growth by inhibiting Wnt/β-catenin both *in vitro* and *in vivo*, and the DUB-independent scaffolding function plays the major role in APC-truncated CRC growth blockade. Clinically, USP10 level significantly correlates with CRC patient survival, Axin1 levels and nuclear β-catenin ratio.

## Discussion

Wnt/β-catenin signaling is an essential pathway that is being actively studied for decades.^1, 2^ For the first time, we identified USP10 as a critical regulator of this pathway and proved that USP10 participates in various Wnt-governed biological processes. In embryonic development and intestinal homeostasis, perturbation of USP10 showed strong Wnt-related phenotypes. And in CRC, USP10 behaves as a strong tumor suppressor both *in vitro* and *in vivo* and shows correlation in the clinical investigations.

As a “star molecule” that was reported to regulate multiple important proteins^21^, the biological significance and multipurpose nature of USP10 are further revealed in this work. However, different from the previous researches, we have discovered a novel mechanism that USP10 regulates Wnt/β-catenin signaling by both classical (DUB activity) and alternative (scaffolding and phase transition) paths. USP10 directly binds to Axin1 through each other’s conserved motifs, which enables the clearance of K48-linked ubiquitination on Axin1 to lengthen the protein lifetime. This binding, on the other hand, occupies the Axin1 intramolecular inhibition site on the DIX domain and potentially allows the “open-state” of Axin1 to further extend for β-catenin entrapment.^11^ Also, the intrinsically-disordered region of USP10 bridges both Axin1 and β-catenin, therefore offers USP10 a scaffold position in the dynamic formation of the destruction complex. As a result, USP10 is crucially for the physical interaction in phase separation-like process and the recruitments of β-catenin. This novel “dual-wielding” mechanism sheds new light on the versatility of USP10 functions and provides new directions for researchers who are interested in DUB studies.

The dual Wnt-regulatory mechanisms of USP10 both rely on Axin1, but can act parallelly to modulate the magnitude of Wnt signaling. We also observed that in different biological processes, the functional significance of each “weapon” was highly context-dependent. For instance, in embryonic development, dorsal-ventral axis formation and patterning primarily require the DUB activity. Both the enzymatic and scaffolding capabilities are involved in intestinal homeostasis, whereas in human CRC the scaffolding capability of USP10 is more predominant. One reason could be that in diverse contexts, the sensitivity of Axin1 stabilization and destruction complex formation are different. This might result in the redundancy of one path and the dearth of the other. Therefore, the dominance of particular activity or the coordination of both in various biological situations requires further scrutiny. Once these processes are fully understood, one could envision that USP10 inhibitor such as Spautin-1, may have therapeutic potentials in particular tissue maintenance, organ regeneration or degenerative disease treatment. And small molecules blocking USP10 PBR binding with Axin1 might have even broader potential applications in Wnt-defective disease therapy.

Many aspects can affect the malignant tumor progression. In our study, USP10 downregulates Wnt/β-catenin signaling to reduce cell stemness and decelerate cell cycle.^1^ Alternatively, USP10 was also reported to inhibits cancer growth by deubiquitinating tumor suppressors including PTEN, AMPK and TP53.^23, 24, 26–28, 54^ So presumably USP10 may affect CRC growth through these proteins. However, the DUB-dead mutant, USP10-CA, which could not deubiquitinate any substrate rather than elevate β-catenin degradation via physical phase separation, showed indistinguishable effect to WT USP10 in Wnt/β-catenin suppression and CRC growth inhibition.

## Supporting information

None

## Significance

To conclude, we have found that USP10 substantially thwarts Wnt/β-catenin signaling in various biological conditions through a novel “dual-regulation” mechanism. This study further discovers a crucial Wnt regulator and offers conceptual breakthrough of DUB enzyme-independent functioning, thus holding great value in basic biomedical research as well as potential for translation.

## Limitations

There are still certain issues for us to dissect in the near future. First, the functions of USP10 in different organ development and homeostasis are not fully studied in this work. Thus, USP10 (conditional) knockout and transgenic animals should be generated to obtain more tissue-specific and physiological results. Second, the relationship between USP10 DUB-dependent and -independent mechanisms in different biological processes needs further scrutiny. It is necessary to better understand the factors in each biological context and then correlate USP10 functionality of individual path.

## Acknowledgement

The authors thank Dr. Chen Ding (Northeastern University) for scientific advice; Dr. Aina He (Shanghai Sixth People’s Hospital) and Dr. Long Zhang (Zhejiang University) for plasmids sharing. R.S. acknowledges the support by the Fundamental Research Fund for the Central Universities, China N182005006 (completed), National Natural Science Foundation of China 31970721 and 81902830, and Liaoning Provincial Talents Project XLYC1807239 (completed). Y.L. acknowledges the National Natural Science Foundation of China 32171248 and 12102142, and the Fundamental Research Funds for the Central Universities, HUST grant 2021GCRC056. Q.W. acknowledges the support of National Natural Science Foundation of China 32025014 and 81921006. L.C. acknowledges the support by the key project of the National Natural Science Foundation of China 82030091.

## Author contributions

R.S. designed the study. Y.W., A.M., J.L., P.L., S.Z., T.T., Z.L., H.Z., L.M., J.L., Q.H., F.Q. Y.L. and R.S. performed experiments and collected and analyzed the data. X.Z (Xinjun Zhang)., M.C., X.H., X.Z (Xi Zhang)., T.F., B.L., L.C., Q.W., Y.L., D.G. and R.S. wrote and revised the manuscript. L.C., Q.W., Y.L. and R.S. oversaw the study. All authors have approved the manuscript.

## STAR Methods

### Key reagents and resource

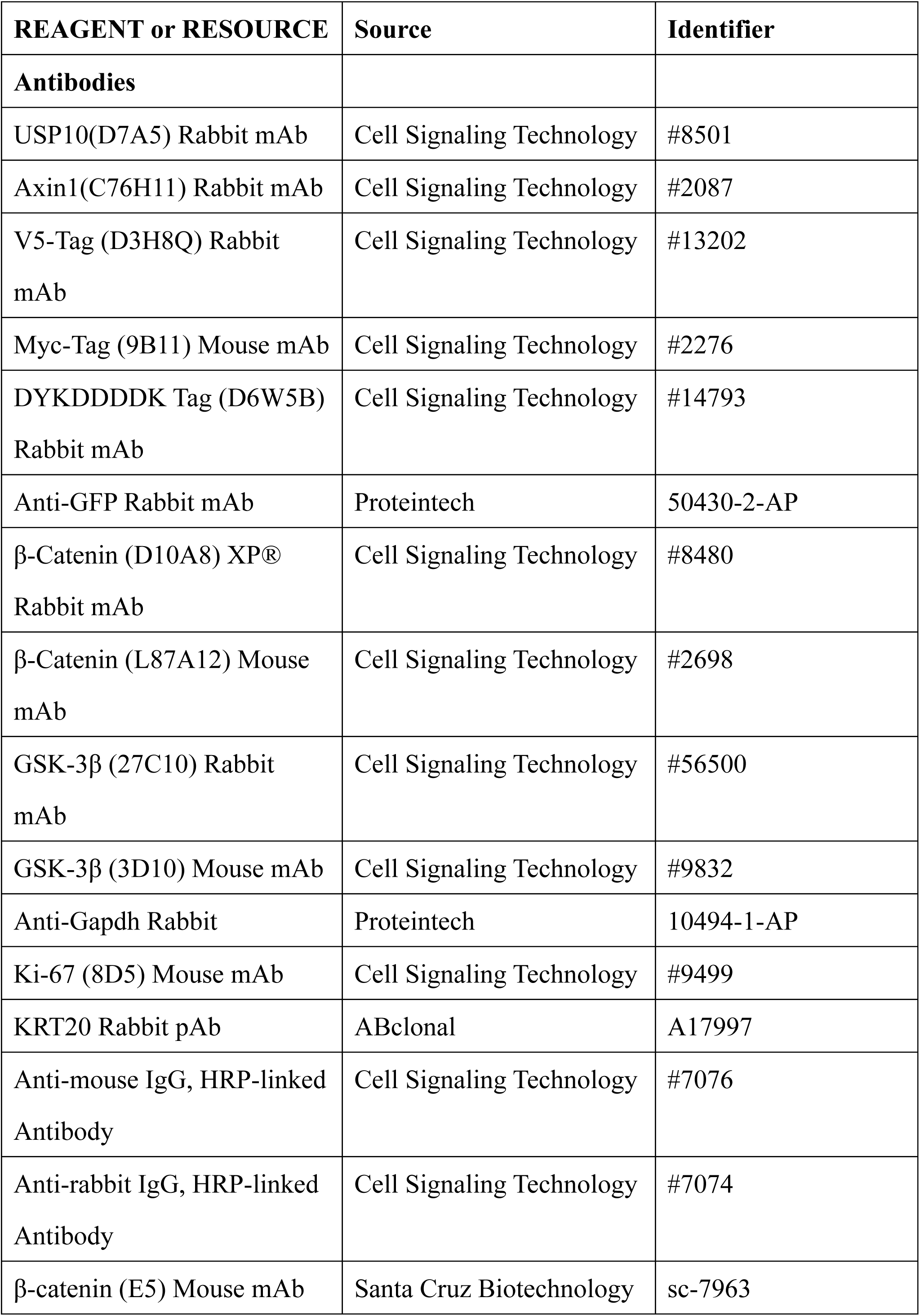
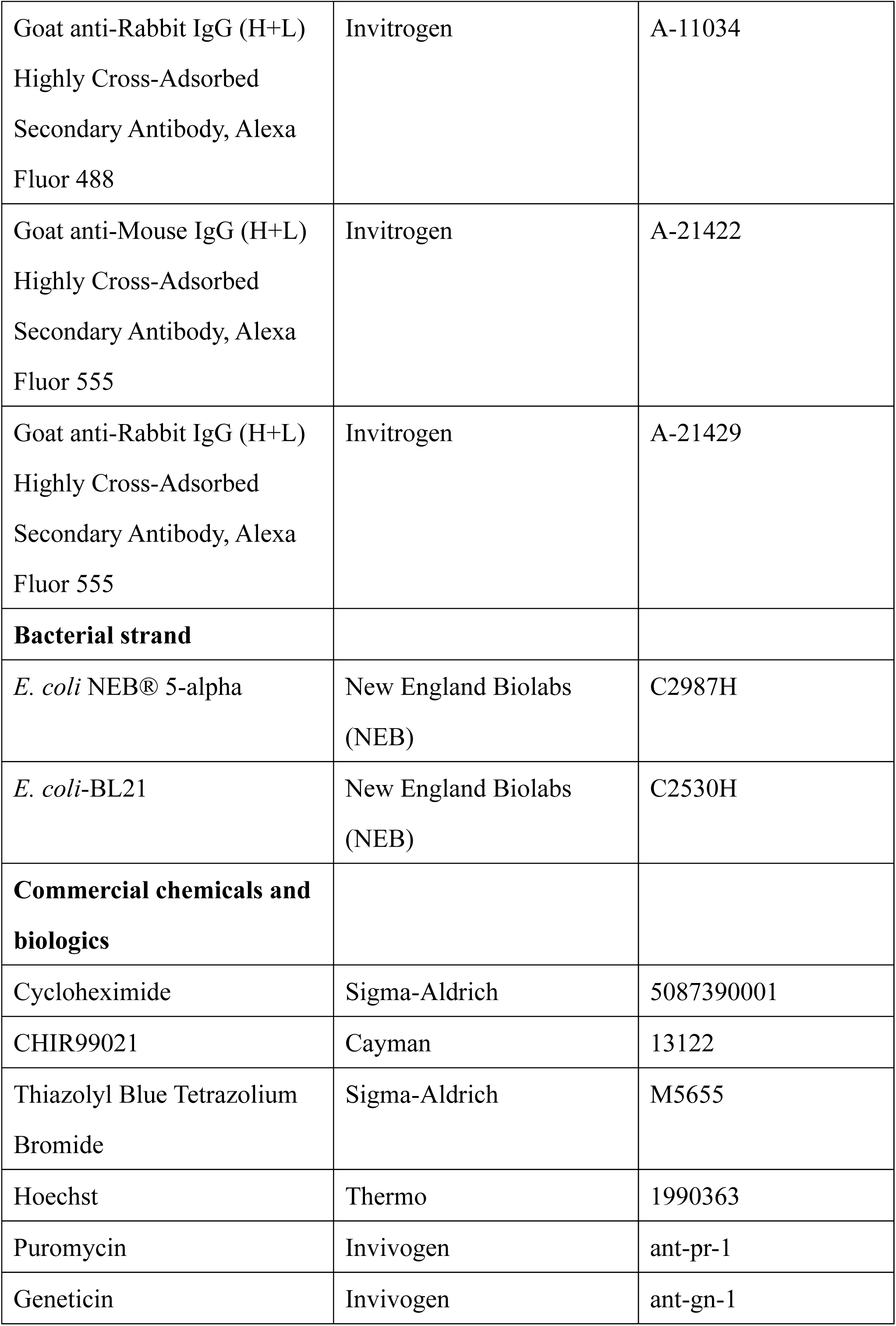
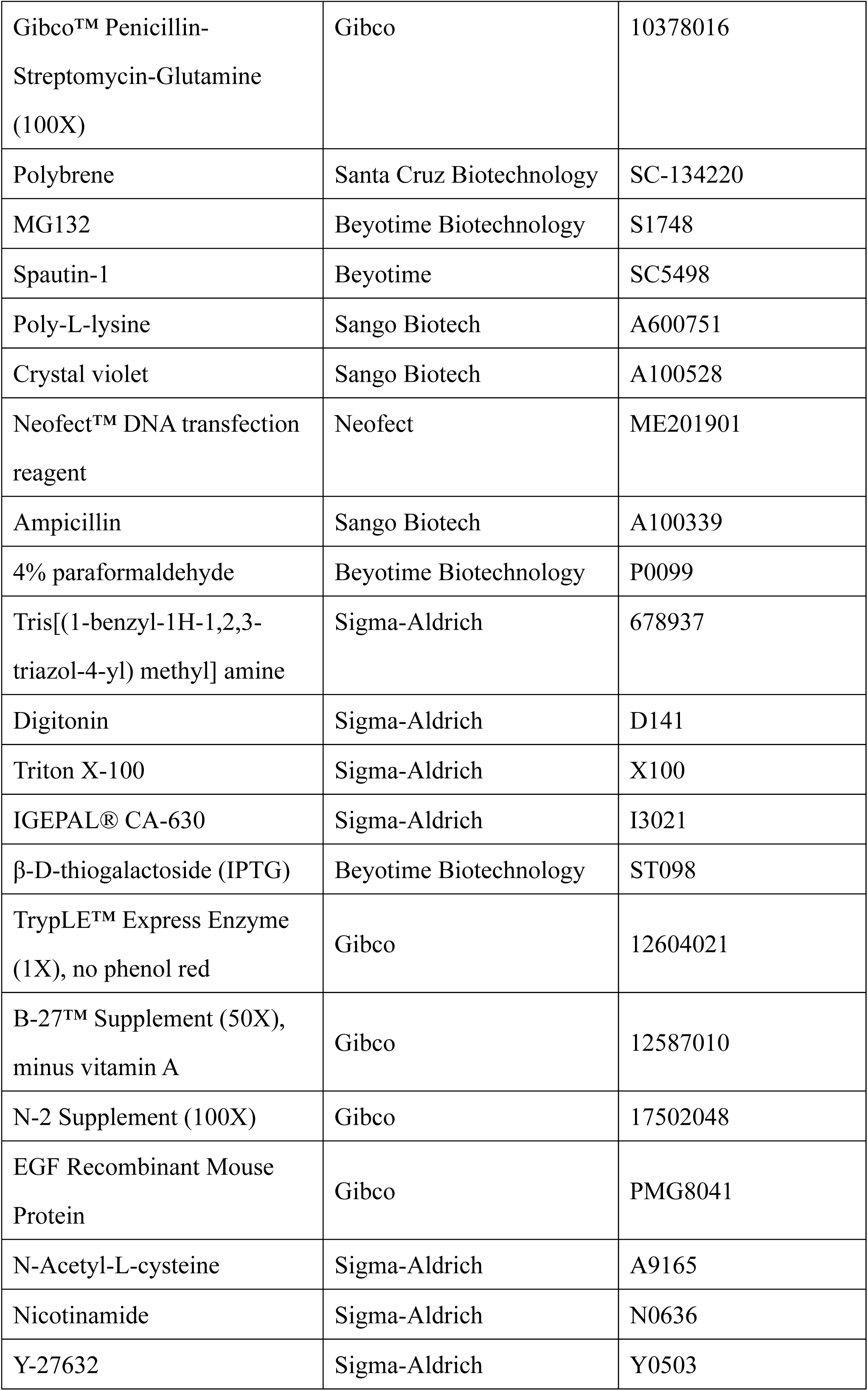
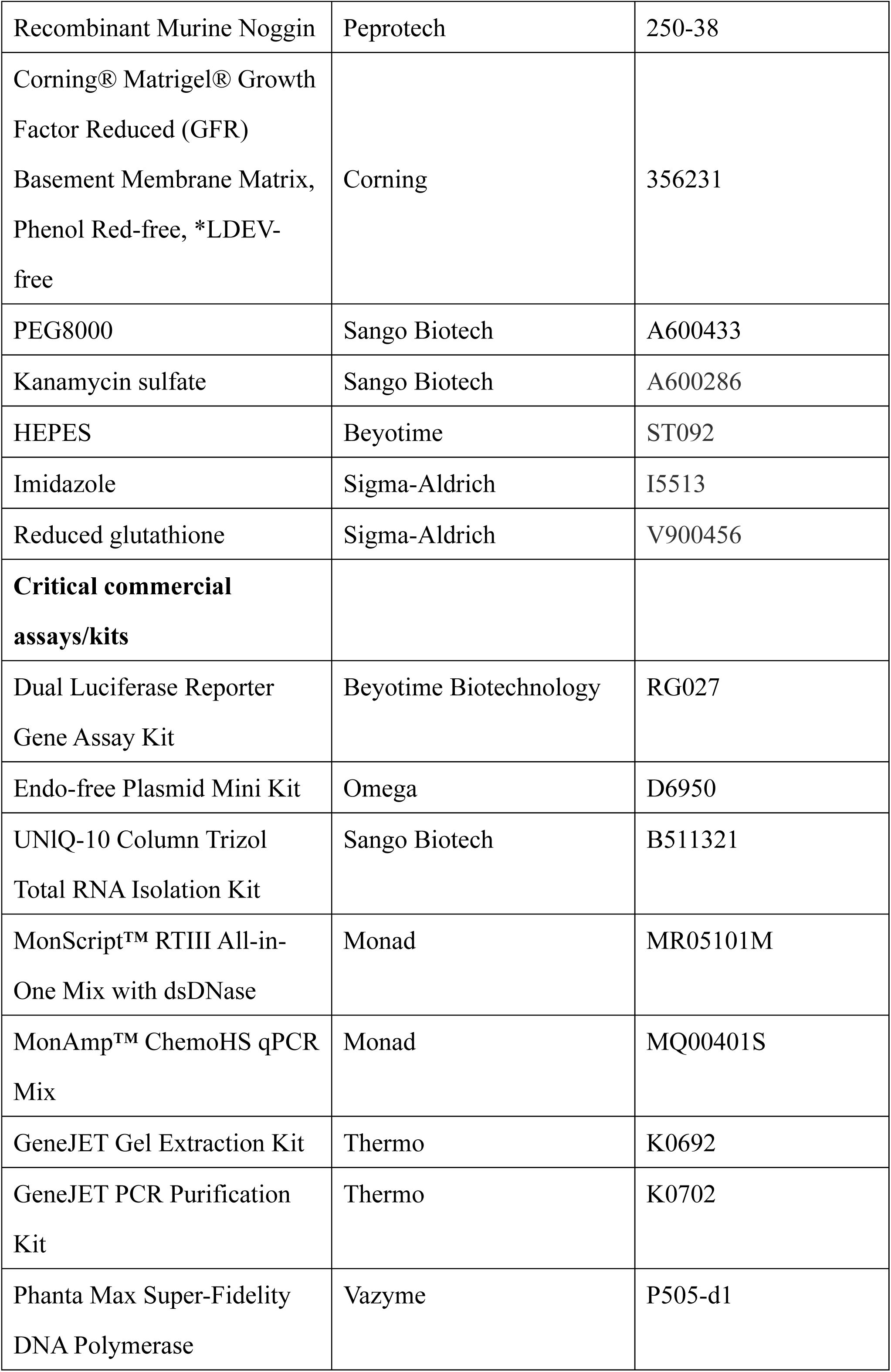
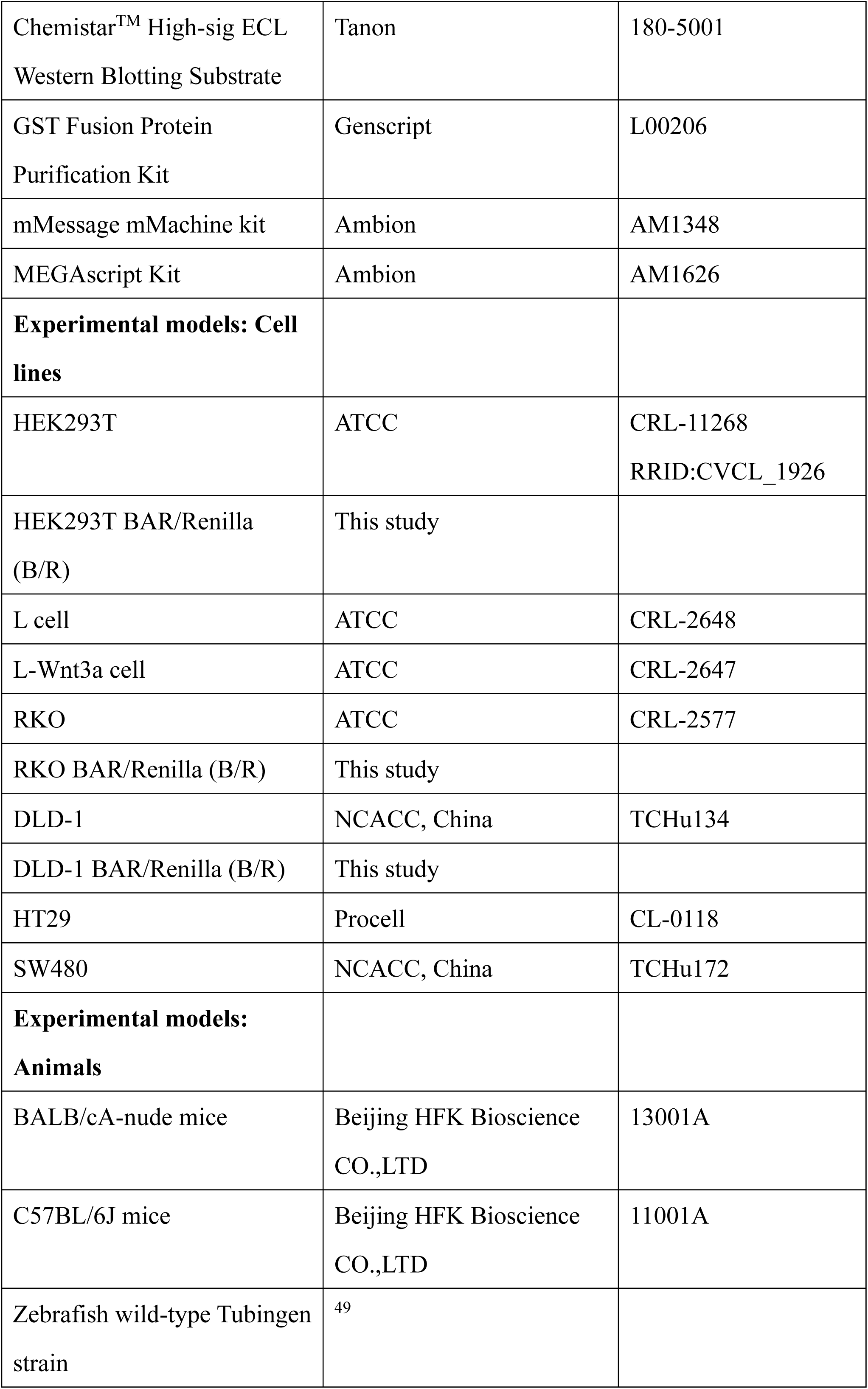
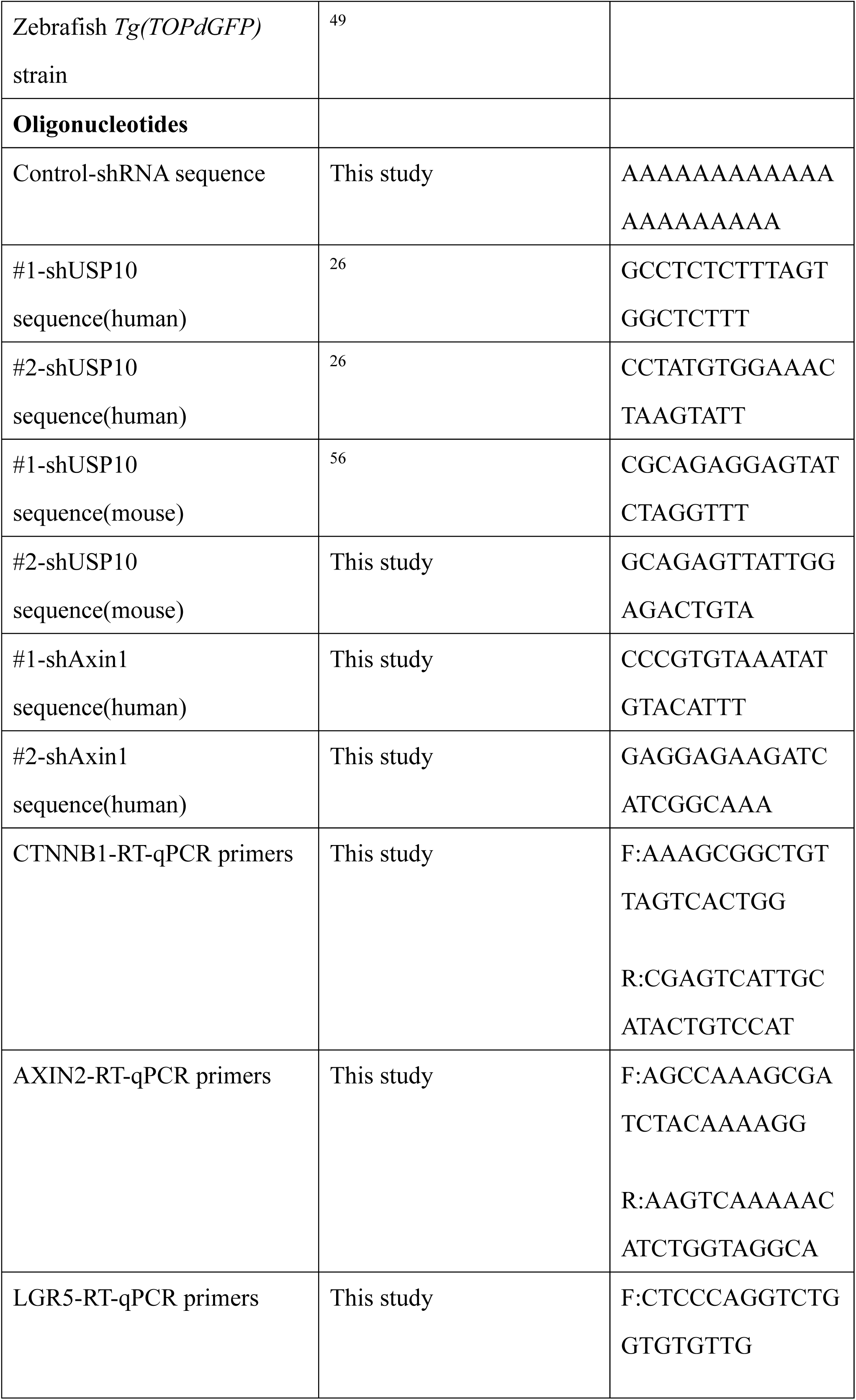
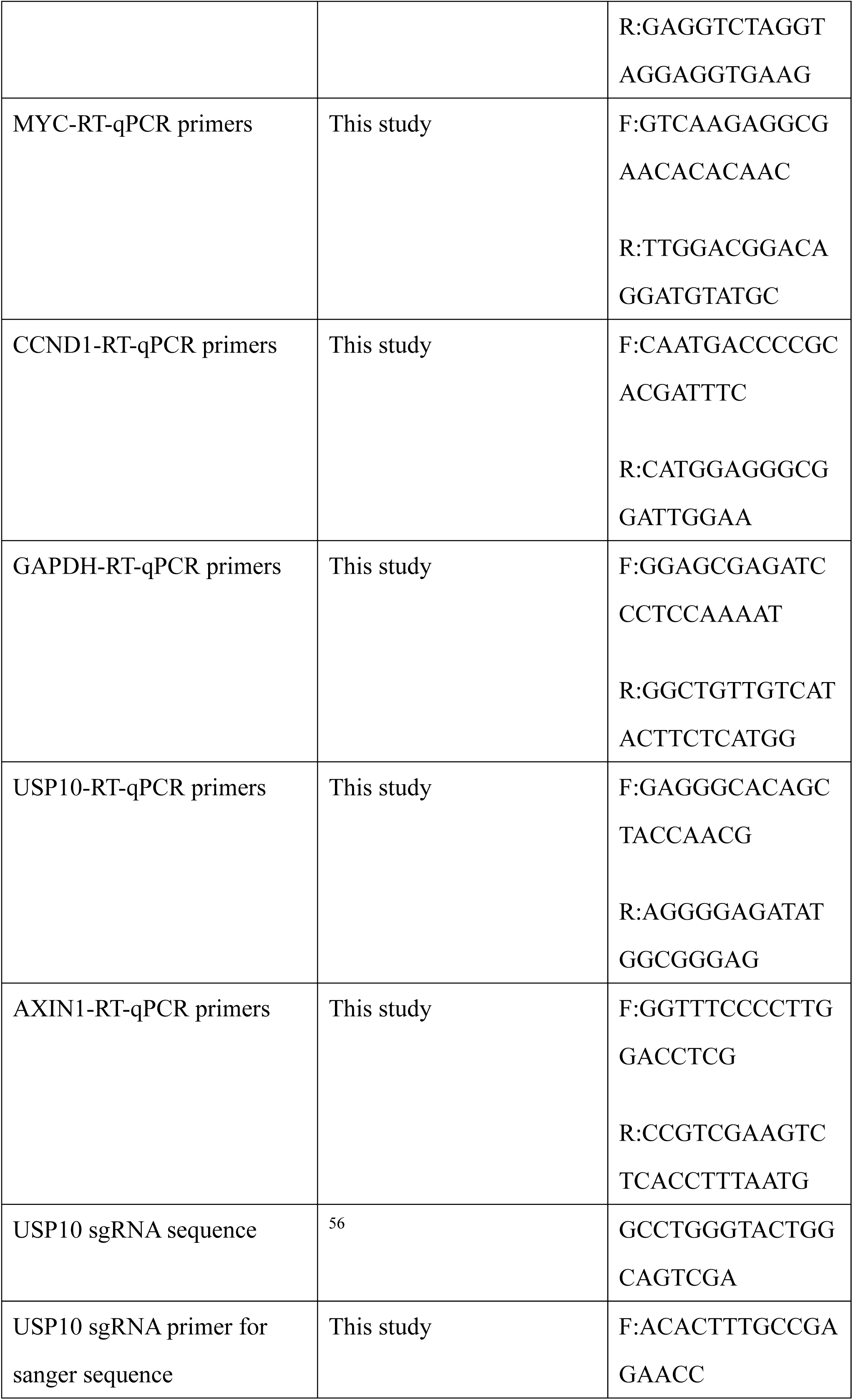
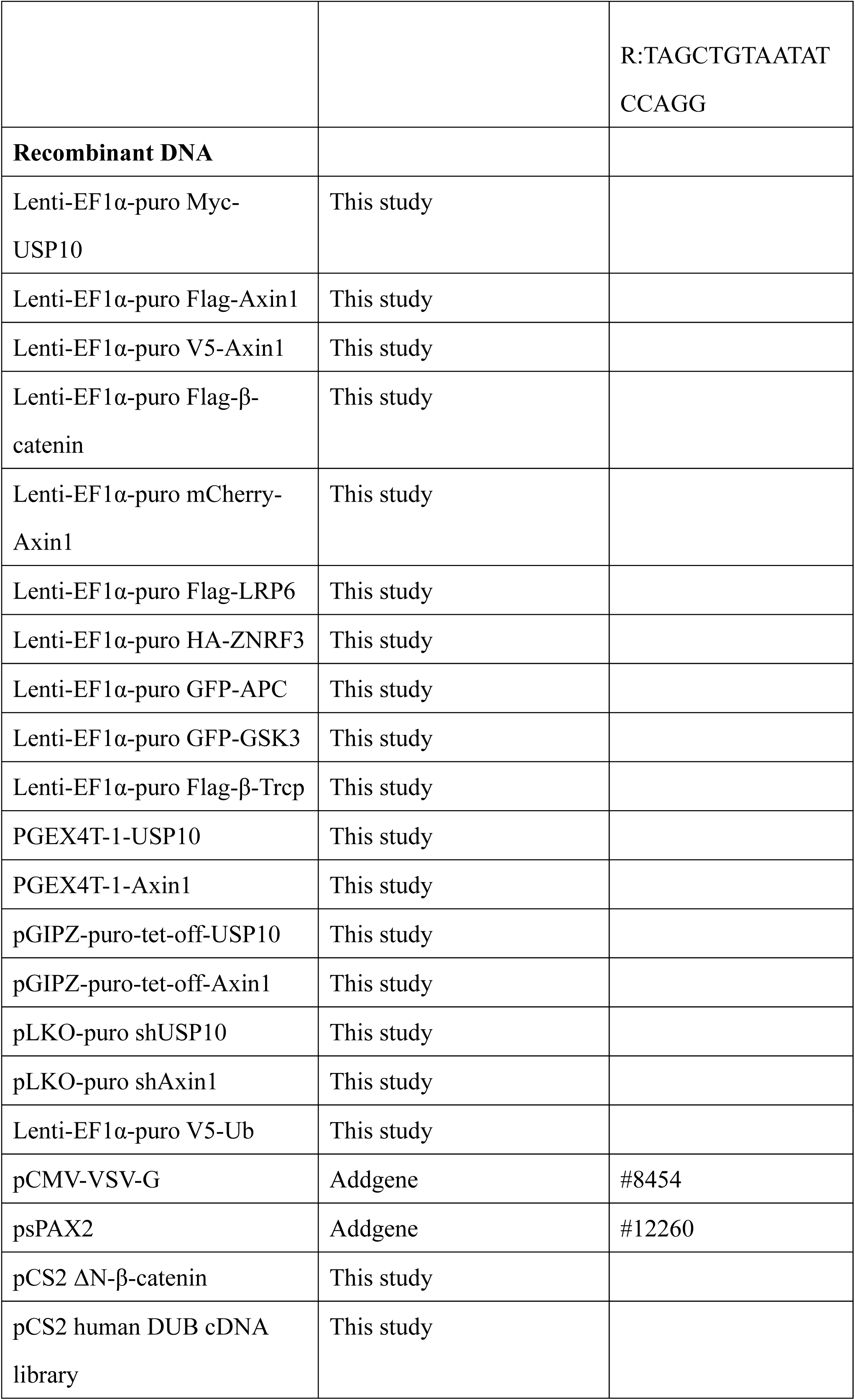
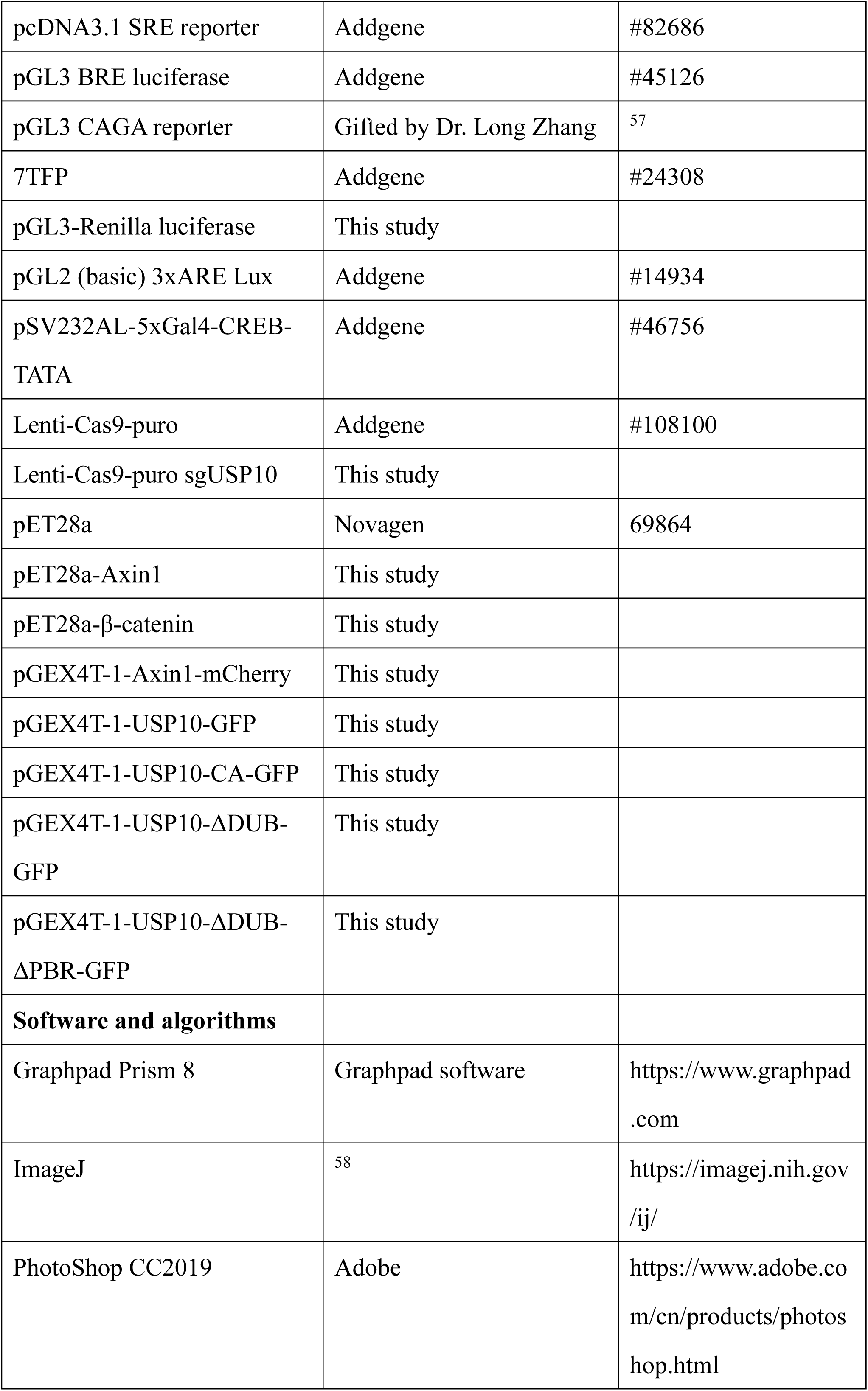
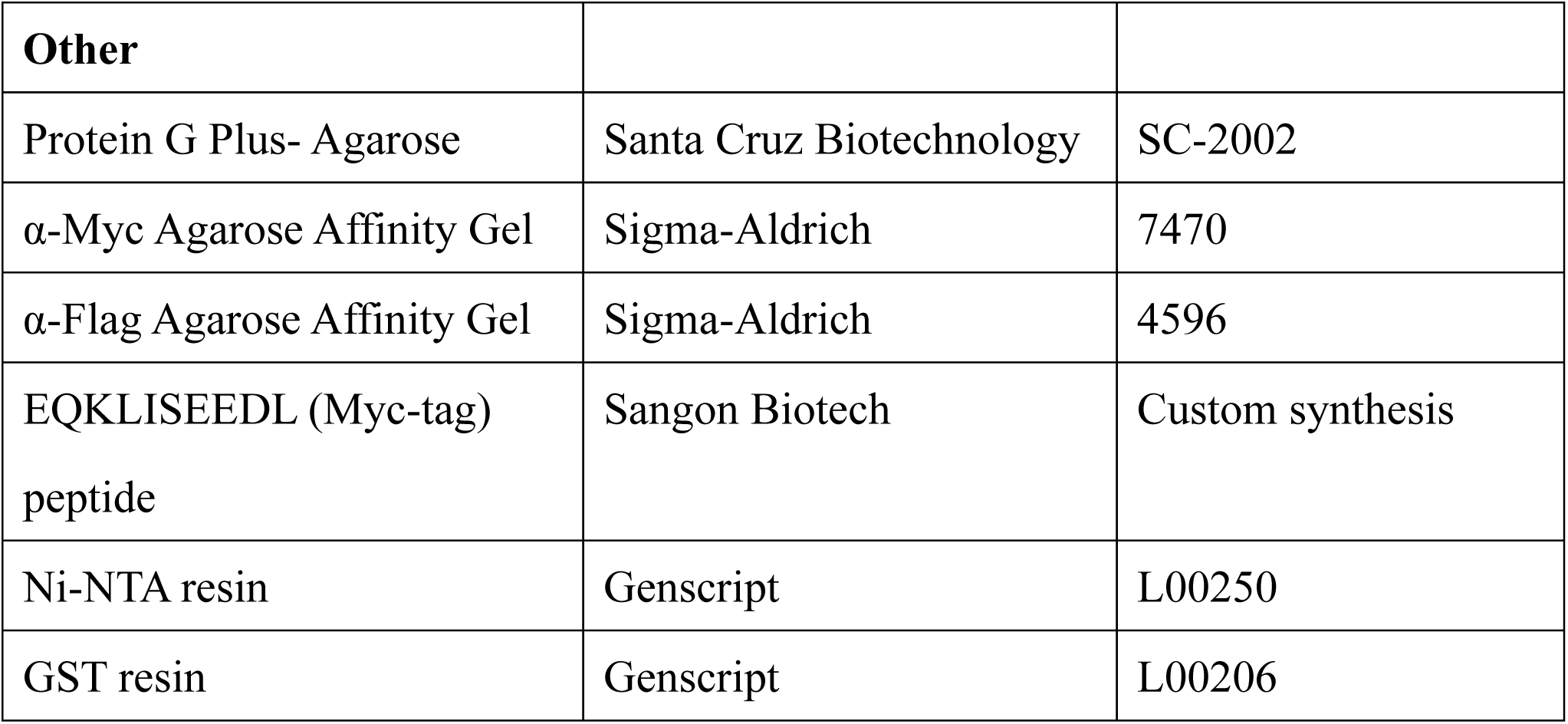

### Human specimen study approval

The tissue microarrays of 92 cases of colon adenocarcinoma (OD-CT-DgCol04) were purchased from Shanghai Outdo Biotech Company, China. The research approach of immunohistochemical detection of protein expression in tissues of colon cancer has been approved by the Ethics Committee of China Medical University (Permission no: [2021] 206).

### Animal work approval

Mice: All mice (BALB/cA-nude and C57BL/6J) were purchased from Beijing HFK Bioscience CO., LTD. Mice maintenance and treatments described were approved by the Research Ethics Committees of the College of Life and Health Sciences of Northeastern University (Approval no. NEU-EC-2021A018S and NEU-EC-2022A019S).

Zebrafish strains: The adult fishes were raised under standard conditions. Embryos were obtained from natural mating, which were grown at 28.5°C in Holtfreter’s solution, and staged according to the morphology as previously described ^59^. Adult wild-type (Tubingen strain) and *Tg(TOPdGFP)* ^49^ were used in this study. The experiments performed were approved by the Animal Care and Use Committee at the Institute of Zoology, Chinese Academy of Sciences (Permission Number: IOZ-13048).

### Subcutaneous tumor transplantation

DLD-1 cells were trypsinized and resuspended in DMEM. Nude mice (Beijing HFK Bioscience CO., LTD) were subcutaneously injected at a density of 1 × 10^6^ DLD-1 cells per site, and the designed cell number and viability were determined using trypan blue.

### Murine intestinal crypt isolation

Mice after anesthesia treatment were sacrificed by cervical dislocation, and 2-3cm portions of the proximal intestine were collected, opened longitudinally and washed with ice-cold PBS. The luminal side was scraped to remove luminal contents and villous structures. After washing again with ice-cold PBS, the intestine was cut into 2-5mm pieces with scissors. The intestinal fragments were then incubated in 2mM EDTA/PBS on ice for 30min. EDTA was then removed, which was followed by 10mL of cold PBS addition for sequel manual suspension for 3min to release the crypt. The supernatant was collected and passed through a 70μm filter (FALCON) to remove tissue debris. After three washes with PBS and centrifugation at 600g to remove tissue debris, the crypts were enriched in the resulting pellet and subsequently embedded in Matrigel gels (Corning).

### Cell lines

This study utilized HEK293T (ATCC), DLD-1 (NCACC), RKO (ATCC), HT29 (Procell), SW480 (NCACC), L (ATCC) and L-Wnt3a (ATCC) cells. All cell lines were maintained in humidified incubators with 5% CO2 at 37°C. HEK293T (parental and genetically modified), DLD-1 (parental and genetically modified), RKO (parental and genetically modified), SW480 (parental and genetically modified), L and L-Wnt3a cells were cultured in DMEM-High Glucose supplemented with 10% FBS (fetal bovine serum) and 100mg/mL of penicillin/streptomycin/glutamine (Gibco). HT29 cells (parental and genetically modified) were cultured in RPMI-1640 medium supplemented with 10% FBS and 100mg/mL of penicillin/streptomycin/glutamine (Gibco).

### Clones and constructs

Expressing Myc-tagged USP10 plasmids (wild-type and mutants), Flag-tagged Axin1 plasmid (wild-type and mutants), Flag -tagged β-catenin plasmid (wild-type and mutants), mCherry-tagged Axin1 plasmids were constructed in customized Lenti-EF1α-puro vector. Plasmids containing shControl, shUSP10 and shAxin1 were constructed in pLKO vector (primer sequence in reagent). Expressing GST-tagged USP10 plasmids (wild-type and mutants) and GST-tagged Axin1 plasmids (wild-type and mutants), GST-tagged USP10-GFP plasmids (wild-type and mutants) and GST-tagged Axin1-mCherry were constructed in pGEX-4T-1 vector. USP10 WT, USP10-CA, Axin1 were constructed in customized pGIPZ-tet-off-puro vector. His6-tagged Axin1 and His6-tagged β-catenin were constructed in pET28a vector. The plasmid containing sgUSP10 was constructed in Lenti-Cas9-puro vector (sequence in the reagent). All plasmids were transformed in *E. coli* NEB^®^5α chain for amplification and extracted by OMEGA Endo-free Plasmid Mini Kit. The concentrations of all plasmids were determined by Thermo Nanodrop 2000.

### Cell culture and transfection

HEK293T, RKO, L and L-Wnt3a cells were purchased from ATCC. DLD-1 and SW480 cells were purchased from the NCACC, China. HT29 were purchased from Procell. HEK293T (parental and genetically modified), RKO (parental and genetically modified), DLD-1 (parental and genetically modified), SW480 (parental and genetically modified), L and L-Wnt3a cells were cultured in DMEM medium supplemented with 10% FBS and 100mg/mL of penicillin/streptomycin/glutamine. HT29 cells (parental and genetically modified) were cultured in RPMI1640 medium with 10% FBS and 100mg/mL penicillin/streptomycin/glutamine.

Transfection was done using Neofect. Transient cDNA transfection was performed using Neofect according to the manufacturer’s recommendations. Plasmids were diluted using DMEM and mixed with Neofect. The complexes were incubated for 20 min at room temperature (RT) and added to HEK293T (parental or modified) cells in growth medium. After 24-48hrs, cells were lysed using Passive Lysis Buffer (25mM Tris-HCl, 150mM NaCl, 0.5% CA630).

### Lentivirus production and infection

For lentivirus production, psPAX2 (for packaging, Addgene) and pCMV-VSV-G (for enveloping, Addgene) and the desired plasmid constructed in the custom Lenti-EF1α-puro vector or Lenti-Cas9-puro sgUSP10 were co-transfected in HEK293T cells at a mass ratio (ng) of 5:1:5. After transfection for 24hrs, the medium was carefully aspirated and replaced with fresh medium to produce virus-containing conditioned medium. Virus can be harvested at 48, 72, and 96hrs post transfection. For lentiviral infection, cell cultures were added to 0.5-1mL of conditioned medium containing virus, 1mL fresh medium and polybrene (Santa Cruz Biotech, 1:1000). After 48hrs, the medium was gently aspirated from the cells and replaced with fresh medium containing appropriate antibiotics for resistance selection.

### Cell line generation

HEK293T BAR/Renilla (B/R) cell line, RKO BAR/Renilla (B/R) cell line and DLD-1 BAR/Renilla (B/R) cell line: HEK293T cells, RKO cells or DLD-1 cells were infected with lentiviruses containing 7TFP and Renilla for 48hrs. After that, the medium was removed and replenished with fresh medium containing puromycin (Invivogen) and geneticin (Invivogen) for selection for 72hrs. Eventually, selected cells were cultured in normal medium and verified by dual luciferase reporter assay. USP10 KO cell lines: The selected sgRNA sequence (GCCTGGGTACTGGCAGTCGA) were cloned into Lenti-Cas9-puro vectors (Addgene). HEK293T or SW480 cells stably expressing Lenti-Cas9-puro sgUSP10 were generated following lentiviral infection and puromycin resistance selection. After reaching confluency, cells were digested and diluted to approximately one cell/well, and seeded into 96-well plates to generate single clone in HEK293T or SW480 cells. Genotypes of single cell clones were determined by both sequencing DNA fragments containing targeted sgRNA regions amplified using sgUSP10 sanger sequence PCR primers and immunoblotting against USP10 antibody (see Reagents or Resource).

### Antibodies and immunoblotting

Antibodies were purchased from different companies (see detail in Reagents and Resources: Antibodies). Cells were lysed by passive lysis buffer (25mM Tris-HCl, 150mM NaCl, 0.5% CA630) containing protease inhibitor cocktail (Roche). The cells were lysed in SDS loading buffer and boiled for 5-10min, followed by 8% or 10% poly-acrylamide gel for SDS-PAGE. For Western blotting, all primary antibodies were used at a 1:1000 dilution and all secondary antibodies were used at a 1:5000 dilution. Chemiluminescent substrate kit was purchased from GE and Tanon. Final quantification of gel intensity was done by ImageJ and plotted in Prism 8.0.

### Immunoprecipitation

For co-immunoprecipitation, total lysates of cells were incubated with α-Flag-agarose or α-Myc-agarose overnight at 4°C. Next day, the resins were washed thoroughly five times with lysis buffer incubated and shaken for 10min at 4°C and resuspended in SDS loading buffer, boiled for 5min, used for the SDS-PAGE and Western blotting.

### Reverse transcription and quantitative real-time PCR

UNlQ-10 Column Trizol Total RNA Isolation Kit (Sango Biotech) was used to extract total RNA from cells and reverse transcribed by MonScript™ RTIII All-in-One Mix with dsDNase (Monad) according to the manufacturer’s protocol. Quantitative RT-PCR (qPCR, for RNA) and PCR (for genomic DNA) were performed using MonAmp™ ChemoHS qPCR Mix (Monad). All primers are designed based on the primer bank of Massachusetts General Hospital (https://pga.mgh.harvard.edu/cgi-bin/primerbank). All experiments were performed in triplicate. The expression values were normalized to those of GAPDH. PCR primer sequences are listed in the “Reagents or Resource”.

### Pulse-chase assay

The cells in 24-well plates (with 70-80% confluency) were treated with cycloheximide (Sigma-Aldrich) 300μg/mL at a range of time points (i.e., 0, 2, 4, 6, 8hrs). Cells were harvested at indicated time points and lysates were prepared and analyzed. Afterwards, the gel intensity was quantified by ImageJ and fitted into first order decay, and protein half-life (t1/2) was calculated according to ln(N)-ln(N0) = -kt linear equation. k=degradation rate constant (min^-1^).

### L conditioned medium (LCM) and L-Wnt3a conditioned medium (Wnt3aCM)

L cells and L-Wnt3a cells were grown in DMEM medium supplemented with 10% FBS. L cells expressing Wnt3a were cultured to 80-90% confluency and collected every 2 days for 6 days. Maximal activity was determined using the TOPFlash assay for subsequent Wnt stimulation experiments. Conditioned medium from L cells collected as control.

### TOPFlash Dual-luciferase reporter assay

For TOPFlash reporter assays, the TOPFlash reporter cell line (HEK293T B/R, cells containing stably expressed superTOPFlash (BAR) and *Renilla* (internal control) vectors) were used. Cells were seeded in 48-well plates and used Neofect for transfect, each set was repeated three times. For experiments requiring Wnt3a stimulation, the medium was changed to 1:1 mixed fresh medium and Wnt3aCM or LCM. After 18 hours of stimulation, dual-luciferase reporter assays were performed using the Dual Luciferase Reporter Assay Kit (Beyotime Biotechnology) according to the manufacturer’s instructions. Plates ready for testing were measured by a Biotek Synergy H1 plate reader. Representative results consist of three (or more) independent experiments.

### Cellular content fractionation

Cell membranes and cytoplasm were separated using digitonin lysis buffer (25mM Tris-HCl, 150mM NaCl, 0.015% digitonin): first, cells were lysed using digitonin lysis buffer for 10min and centrifuged at 6124xg for 2min at 4℃, and the supernatant (cytoplasmic fraction) collected, resuspended in SDS loading buffer, boiled for 5min, used for the SDS-PAGE and Western blotting.

### *In vivo* and *in vitro* deubiquitination assay

For *in vivo* deubiquitination, V5-tagged Ub plasmid and other needed plasmids were transfected in HEK293T cells, and the cells were then treated overnight with 10μM proteasome inhibitor MG132 (Beyotime Biotechnology). These cells were lysed with cell lysis buffer after 48hrs and incubated with a certain amount of α-Flag-agarose overnight at 4℃, and analyze by Western blotting.

For *in vitro* deubiquitination experiments, HEK293T cells were co-transfected with Flag-Axin1 and V5-Ub treated with 10μM MG132 overnight. After 48hrs, cells were lysed and 200μL was left as input, and the remaining whole cell lysate was incubated with a certain amount of Flag beads overnight at 4℃. After binding, the beads were washed with lysis buffer for later use. Meanwhile Myc-USP10 WT and Myc-USP10 CA were expressed in HEK293T cells and purified using α-Myc-agarose. Purified proteins were eluted with 100ng/mL Myc-tag peptide. Eluted USP10 WT or USP10 CA proteins were then incubated with prepared Flag-Axin1 beads in deubiquitination buffer (50mM Tris-HCl, pH 7.4, 1mM MgCl2 and 1mM DTT) for 2hrs at 37℃. The beads (Flag-Axin1) were washed five times by TBS with sufficient shaking during each interval and then analyzed by Western blotting.

### Pull-down assay

Preparation of GST fusion protein: Recombinant plasmids with the target gene in pGEX-4T-1 were transformed to *E. coli* BL21 competent cells. Protein expression was induced by addition of IPTG (Beyotime Biotechnology) and shaken at 3xg for 6-8hrs at 18℃. The bacterial solution was then collected, centrifuged, and the pellet was resuspended in 20 mL of ice lysis buffer (1mM PMSF, Tris, NaCl, and glycerol), and sonicated by SCIENTZ JY92-IIN sonicator on ice at 40% power for 5s each time at 5s intervals for 20min. The bacterial solution after sonication was centrifuged at 4℃ for 10min. Transfer the supernatant to a 50mL centrifuge tube. An appropriate amount of GST resins was added to the lysis buffer and incubated for 2-3hrs at 4℃ with rotation. The binding beads were washed three times in ice-cold TBS and brought to volume with 1mL TBS (1mM PMSF) for GST-pull down experiments.

GST-pull down: GST-pull down analysis was performed to collect HEK293T cell lysates transferred with the target protein. A small amount of supernatant was obtained as input control, and the remaining lysate was added with beads coupled with GST-fusion expression protein and incubated overnight at 4°C. Beads were washed three times with lysis buffer, the supernatant was aspirated, and an appropriate amount of loading buffer, resuspended in SDS loading buffer, boiled for 5min, used for the SDS-PAGE and Western blotting.

Preparation of His6-tagged protein: The general protocol was the similar with GST-tagged protein preparation. The Ni-NTA resins were used to enrich His6-tagged protein and imidazole was used to elute. Imidazole was removed by dialysis at 4°C overnight.

### Whole-mount *in situ* hybridization

Digoxigenin-UTP-labeled antisense RNA probes were synthesized in vitro using the MEGAscript Kit (Ambion) according to the manufacturer’s instructions. Whole-mount *in situ* hybridization (WISH) with the RNA probes was performed as previously described methods.^60^

### RNA microinjections

Capped mRNAs were synthesized in vitro for USP10, GFP from the corresponding linearized plasmids using the mMessage mMachine kit (Ambion) according to the manufacturer’s instructions. The mRNA was injected into the embryos at 1-cell stage with a concentration of 500ng/μL or co-injected with 100ng/μL ΔN-β-catenin mRNA, 1nL per embryo.

### Chemical treatments

The USP10 inhibitor, Spautin-1(Beyotime), was dissolved in DMSO with a concentration of 10mM as stock solution. HEK293T were treated with the inhibitor Spautin-1 at a final concentration of 10μM. Embryos were treated with the inhibitor Spautin-1 at a final concentration of 5μM or 10μM (diluted in Holtfreter’s solution) form 8-cell stage. Intestinal organoids were treated with the inhibitor Spautin-1 at a final concentration of 20μM, and then harvested at the indicated stage for further analysis.

### Mouse intestinal organoids culture

The general process was described in ^50^. Freshly isolated mouse crypts or single cells from dissociated mouse intestinal stem cell (ISC) colonies were embedded in Matrigel gels (Corning), which were cast into 20μL droplets at the bottom of wells in 96-well plate. Following polymerization (10min, 37°C), the gels were overlaid with 150μL of ISC expansion medium (Advanced DMEM/F12 containing Glutamax, HEPES, penicillin-streptomycin, B27, N2 (Gibco), 1μM N-acetylcysteine (Sigma) and 10mM Nicotinamide (Sigma), supplemented with growth factors, including EGF (50ng/mL; Gibco), Wnt, Noggin, R-spondin1 (produced in-house), and small molecules, including CHIR99021 (5μM; Sigma) and Spautin-1 (20μM; Beyotime). The organoids were infected by lentivirus for 72hrs. The specific protocols were previously described.^61^

### Immunofluorescence analysis

Mouse intestinal organoids embedded in Matrigel gels were fixed with 4% paraformaldehyde (PFA) in PBS (30min, room temperature). The fixation process typically led to complete degradation of the Matrigel. Suspended tissues were collected and centrifuged (61xg, 5min) to remove the PFA, washed with ultra-pure water and pelleted. Following resuspension in water, the organoids were spread on glass slides and allowed to attach by drying. Attached organoids were rehydrated with PBS. Following fixation, organoids spreaded on glass were permeabilized with 0.2% Triton X-100 in PBS (1h, room temperature) and blocked (10% goat serum in PBS containing 0.01% Triton X-100) for at least 3hrs. Samples were subsequently incubated overnight at 4°C with primary antibodies against Ki67 (1:50; CST #9499) and KRT20 (1:50; ABclonal A17997) diluted in blocking buffer. After washing with PBS for at least 3hrs, samples were incubated 2hrs at 4°C with secondary antibody (1:1000 in blocking solution; Invitrogen). Following extensive washing, stained organoids were imaged in confocal microscope (DM6000CS, Leica).

SW480 (parental or modified) cells were grown on glass coverslips. After growing to 60% confluency, cells were washed three times with PBS and fixed in 4% paraformaldehyde (Sigma), followed by blocking in 5% BSA with 0.2% Triton X-100 for 30min and incubation with primary antibody Axin1 (#2087, 1:200, Cell Signaling Technology) overnight. After that, secondary antibody (A-21429,1:500, Invitrogen) for 40min and nuclei were stained with Hoechst/DAPI (Thermo). Finally, samples were visualized using a confocal fluorescent microscope (DM6000CS, Leica).

### *In vitro* droplet formation assays

Axin1-mCherry, USP10 WT-GFP, USP10 CA-GFP, USP10-ΔDUB-GFP and USP10-ΔDUB-ΔPBR-GFP were transformed into *E. coli* BL21 strand for IPTG-induced expression (see Pull-down assay), followed by GST-resin packed column for purification. Reduced glutathione was used to elute the protein. After overnight dialysis at 4°C, the purified proteins were stored in phase separation buffer (20mM HEPES, 1M NaCl, pH 7.4). NaCl concentration was adjusted to the indicated concentration (150mM, pH 7.4) with buffer containing 20mM HEPES at the time of used. Mixtures were immediately treated with PEG8000 (Differences in the nature of phase separation between proteins determine the critical concentration at which droplets are formed).10μL of the reaction mixture was prepared; the concentration of NaCl was further adjusted to 150mM NaCl, and the mixture was dropped onto a glass slide and covered with a coverslip at the above concentration, allowed to stand for a few minutes, incubated, and then examined for droplet formation. For imaging, droplets were viewed on glass slides or in glass-bottom cell culture dishes for droplet and fluorescence imaging using ZOE^TM^ Fluorescent Cell Imager (BIO-RAD).

### Methyl thiazolyl tetrazolium (MTT) assay for cell proliferation

Cells were seeded on 96-well plates at a density of 1000 cells per well in triplicate, incubated for 1-10 days, and cell density was measured on days 2, 4, 6, 8, and 10 after seeding. At each time point, 10µL of MTT (thiazolyl blue tetrazolium, from Sigma) was added to each well at a final concentration of 0.5mg/mL, and plates were incubated for an additional 4hrs at 37°C. After incubation, all MTT was removed and 100μL DMSO was added to each well. Plates ready for testing were measured after 10 minutes using a Biotek Synergy H1 plate reader at OD490. Growth curves were plotted on a daily basis based on OD490 values. Statistical analysis was done by two-way analysis of variance in Prism 8.0.

### Colony formation assay

At a density of 1000 cells /well, the DLD-1 cells were seeded in 6-well plates, and then the cells were cultured in a 37℃ incubator for 14 days to allow colony formation. After the size and number of clones grew to a certain extent, the cells were washed and treated with PBS, fixed with 4% paraformaldehyde for 30min, stained with 0.5% crystal violet for 20min, washed with ddH2O several times and photographed with a digital camera. All assays were performed in triplicate.

### Cell migration assays

Cells were seeded into six-well plates. After reaching 95-100% cell density, 10μL plastic pipette tips were then used to generate scratches in each well. CRC cells were washed with PBS and then maintained in the medium containing 1% FBS. Wound margins were photographed at 0, 24 and 48hrs. Cell migration ability was assessed by measuring the distance between the advancing edges of cells in the microscopic field at each time point. The formula was as follows: 24hrs migration% = (0hr width - 24hrs width of wound)/(0hr width of wound), 48hrs migration% = (0hr width - 48hrs width of wound)/(0hr width of wound). All of the experiments were repeated three times.

### Growth of cells in athymic nude mouse and tumor size determination

DLD-1 cells were trypsinized, washed and resuspended in 0.1mL DMEM medium without serum. Eighty 6-week-old female athymic mice (Beijing HFK Biotechnology Co., Ltd.) were randomly divided into 8 groups (10 mice/group), and then the designated cells (1 × 10^6^) were subcutaneously injected into the plate. Tumor size was determined every 2-3 days. Tumor volume was measured, and tumor volume was calculated using each formula: ½ (Length × Width^2^). Mice were then sacrificed and tumors were excised, weighed and evaluated by immunohistochemistry.

### Immunohistochemistry

Mouse immunohistochemistry tissue samples were fixed in 10% formalin, embedded in paraffin, sectioned, deparaffinized, treated with 3% hydrogen peroxide for 10min, submerging in citric acid (pH 6.0) and microwaved for antigen retrieval, and cooled at room temperature (RT) and incubated in normal goat serum for 1hr to block non-specific binding, then incubated with hematoxylin-eosin staining for 10min or overnight at 4℃ using the following primary antibodies: β-catenin (1:200, #8480, Cell Signaling Technology), Ki67 (1:400, #9449, Cell Signaling Technology). The staining was examined using HRP Envision Systems (DAB Kit, MXB Biotechnologies) and analyzed using a dissecting microscope (Leica DM4000). The human tissue microarrays were purchased from Shanghai Outdo Biotech Company, China. After deparaffinizing in xylene and rehydrating in graded ethanol, tissue microarrays were immersed in citrate buffer (USP10, Axin1) or EDTA (β-catenin) for antigen retrieval. Endogenous peroxidase was quenched using 3% hydrogen peroxide for 30min. To decrease the nonspecific staining, 10% normal goat serum was subsequently used to block tissue collagen for 30min. Tissue sections were then incubated with antibody anti-USP10 (#8501, 1:100, Cell Signaling Technology), anti-Axin1 (#2087, 1:100, Cell Signaling Technology) or anti-β-catenin (sc-7963, 1:500, Sigma) for 90min at room temperature (24-27°C) at 4°C overnight. After that, biotinylated secondary antibody and streptavidin-biotin peroxidase were used to incubate tissue sections for 10min each in turn. Slides were stained with DAB chromogenic reagent for 60s, afterwards counterstained with hematoxylin. UltraSensitiveTM SPIHC Kit (KIT-9720, Maixin Inc., Fujian, China) were used in this experiment.

The stained sections were reviewed and scored by two investigators independently and the different scoring was resolved after discussion. We adopted a semi-quantitative scoring method to assess the expression of certain protein. The staining intensity was divided into 0 (no staining), 1 (weak staining), 2 (moderate) and 3 (strong). The percentage of cells stained was categorized as 0 (0-5%), 1 (6-25%), 2 (26-50%), 3 (51-75%), and 4 (76%-100%). The final scores were generated by multiplying the staining intensity by percentage of cells.

### Confocal laser scanning microscopy

All plasmid transfection fluorescence images were conducted on Olympus spinning disk confocal microscope with an IX83 fully motorized inverted microscope and a 100XO UPLSAPO oil lens (numerical aperture 1.42) (Olympus). mCherry was excited at 594 nm and detected at 610-710nm.

### Fluorescence recovery after photobleaching (FRAP) experiments

Cells were firstly transfected with the plasmids of interest. After 2 days of transfection and expression of the target proteins, the FRAP experiments of these cells were conducted using a Fluoview FV3000 confocal laser scanning microscope with a 100XO UPLSAPO oil lens (numerical aperture 1.4) (Olympus). A granule of the proteins of interests or a small region of the granule was selected and bleached using a 594nm argon laser at 100% intensity in five repeats with a dwell time of 50ms. Time-lapse images were recorded with an internal of 0.09ms between each frame. FRAP analysis was completed using the ImageJ FRAP Profiler plugin (from the Hardin lab: https://worms.zoology.wisc.edu/research/4d/4d.html#frap). The mobile fraction (fm) and the immobile fraction (fi) was calculated by the following equations:

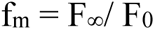

where F∞ is the fluorescence intensity after full recovery, and F0 is the fluorescence intensity before photobleaching.

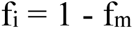

FRAP curve was plotted with Prism 8 software, and *t1/2* was calculated by fitting the curve with exponential decay function.

### Quantification and statistics

P-values were determined with Student’s t-test, One-way or Two-way ANOVA with post hoc Tukey’s HSD test between comparator groups using GraphPad Prism software based on the individual mathematic model of each data set.

WB stripe quantification was performed by Image J:

Using Image J→ Analyze→Gels→Plot Lanes to analyze the Western Blot results. If statistical analysis is required, three independent experiments were performed and quantified. Unpaired two-tailed Student’s t-test was used to determine statistical significance.

For the quantification of Axin1 puncta *in vitro*, maximum projections of 10 z-stack images (0.9µm apart) were manually generated. For the quantification of Axin1 puncta *in vivo*, all images presented in the paper were z-stack maximum projections using a step size of 0.27µm, spanning the entire depth of the nucleus. Puncta number and diameter were analyzed using the software ImageJ. Unpaired two-tailed Student’s t-test was used to determine statistical significance.

TOPFlash assay: After the values were obtained according to the TOPFlash Dual-luciferase reporter assay method, the fluorescence values of Firefly were divided by the corresponding fluorescence values of Renilla (internal control). The values of EV + LCM were used as control, and all the values were divided by the values of EV + LCM for normalization, and the obtained results were used for image drawing.

Experimental results are shown as the mean ± SD, n=3 replicates, one-way ANOVA or two-tailed Student’s t-test was performed for statistical analysis.

MTT assay: The OD490 values at each time point was used as the result for image drawing. Experimental results are shown as the mean ± SD, n=3 replicates, two-way ANOVA was performed for statistical analysis.

Quantitative real-time PCR: The gene expression values were normalized to those of GAPDH. And data processing was performed using the 2^-ΔΔCt^ method, the results were calculated for image drawing. Experimental results are shown as the mean ± SD, n=3 replicates, two-tailed Student’s t-test was performed for statistical analysis.

Tumor volumes: Tumor volume was calculated using the formula: ½ (Length × Width^2^). Images were drawn after obtaining the tumor volumes in combination with the corresponding time points. Experimental results are shown as the mean ± SD, n=10 for each group, two-way ANOVA was performed for statistical analysis.

Tumor weight: The tumor weight was measured after sacrificing the nude mice and dissecting the tumor. Images were drawn based on the weight results. Experimental results are shown as the mean ± SD, n=10 for each group, two-tailed Student’s t-test was performed for statistical analysis.

Endogenous Axin1 puncta in SW480 cells: each group selects three different visual fields (25um), and counts the cells through ImageJ. Experimental results are shown as the mean ± SD, and n was labelled within the figures. Unpaired two-tailed Student’s t-test was used to determine statistical significance.

*In vitro* droplet formation: puncta number and diameter of Axin1 were analyzed using the software ImageJ. Experimental results are shown as the mean ± SD, and n was labelled within the figures. Unpaired two-tailed Student’s t-test was used to determine statistical significance.

Colony number: Image J→ Type:8-bit→Adjust:Threshold→Analyze:Analyze Particles to analyze the colony formation results.^62^ Experimental results are shown as the mean ± SD. Unpaired two-tailed Student’s t-test was used to determine statistical significance.

## Reference

1. MacDonald, B. T., Tamai, K. & He, X. Wnt/beta-catenin signaling: components, mechanisms, and diseases. Developmental cell 17, 9–26, doi:10.1016/j.devcel.2009.06.016 (2009).

2. Nusse, R. & Clevers, H. Wnt/beta-Catenin Signaling, Disease, and Emerging Therapeutic Modalities. Cell 169, 985–999, doi:10.1016/j.cell.2017.05.016 (2017).

3. Storm, E. E. et al. Targeting PTPRK-RSPO3 colon tumours promotes differentiation and loss of stem-cell function. Nature 529, 97–100, doi:10.1038/nature16466 (2016).

4. Bendell, J. et al. Initial results from a phase 1a/b study of OMP-131R10, a first-in-class anti-RSPO3 antibody, in advanced solid tumors and previously treated metastatic colorectal cancer (CRC). Eur J Cancer 69, S29–S30, doi:Doi 10.1016/S0959-8049(16)32668-5 (2016).

5. Chartier, C. et al. Therapeutic Targeting of Tumor-Derived R-Spondin Attenuates beta-Catenin Signaling and Tumorigenesis in Multiple Cancer Types. Cancer research 76, 713–723, doi:10.1158/0008-5472.CAN-15-0561 (2016).

6. Kakugawa, S. et al. Notum deacylates Wnt proteins to suppress signalling activity. Nature 519, 187–192, doi:10.1038/nature14259 (2015).

7. Zhang, X. et al. Notum is required for neural and head induction via Wnt deacylation, oxidation, and inactivation. Developmental cell 32, 719–730, doi:10.1016/j.devcel.2015.02.014 (2015).

8. Flanagan, D. J. et al. NOTUM from Apc-mutant cells biases clonal competition to initiate cancer. Nature 594, 430–435, doi:10.1038/s41586-021-03525-z (2021).

9. Ter Steege, E. J. & Bakker, E. R. M. The role of R-spondin proteins in cancer biology. Oncogene 40, 6469–6478, doi:10.1038/s41388-021-02059-y (2021).

10. Nong, J. et al. Phase separation of Axin organizes the beta-catenin destruction complex. The Journal of cell biology 220, doi:10.1083/jcb.202012112 (2021).

11. Kim, S. E. et al. Wnt stabilization of beta-catenin reveals principles for morphogen receptor-scaffold assemblies. Science 340, 867–870, doi:10.1126/science.1232389 (2013).

12. Huang, S. M. et al. Tankyrase inhibition stabilizes axin and antagonizes Wnt signalling. Nature 461, 614–620, doi:10.1038/nature08356 (2009).

13. Zhang, Y. et al. RNF146 is a poly(ADP-ribose)-directed E3 ligase that regulates axin degradation and Wnt signalling. Nature cell biology 13, 623–629, doi:10.1038/ncb2222 (2011).

14. Callow, M. G. et al. Ubiquitin ligase RNF146 regulates tankyrase and Axin to promote Wnt signaling. PloS one 6, e22595, doi:10.1371/journal.pone.0022595 (2011).

15. Ji, L. et al. USP7 inhibits Wnt/beta-catenin signaling through promoting stabilization of Axin. Nature communications 10, doi:ARTN 4184 10.1038/s41467-019-12143-3 (2019).

16. Ji, L. et al. The SIAH E3 ubiquitin ligases promote Wnt/beta-catenin signaling through mediating Wnt-induced Axin degradation. Genes Dev 31, 904–915, doi:10.1101/gad.300053.117 (2017).

17. Lui, T. T. et al. The ubiquitin-specific protease USP34 regulates axin stability and Wnt/beta-catenin signaling. Molecular and cellular biology 31, 2053–2065, doi:10.1128/MCB.01094-10 (2011).

18. Cha, B. et al. Methylation by protein arginine methyltransferase 1 increases stability of Axin, a negative regulator of Wnt signaling. Oncogene 30, 2379–2389, doi:10.1038/onc.2010.610 (2011).

19. Kim, S. & Jho, E. H. The protein stability of Axin, a negative regulator of Wnt signaling, is regulated by Smad ubiquitination regulatory factor 2 (Smurf2). The Journal of biological chemistry 285, 36420–36426, doi:10.1074/jbc.M110.137471 (2010).

20. Swatek, K. N. & Komander, D. Ubiquitin modifications. Cell research 26, 399–422, doi:10.1038/cr.2016.39 (2016).

21. Bhattacharya, U., Neizer-Ashun, F., Mukherjee, P. & Bhattacharya, R. When the chains do not break: the role of USP10 in physiology and pathology. Cell Death Dis 11, 1033, doi:10.1038/s41419-020-03246-7 (2020).

22. Weisberg, E. L. et al. Inhibition of USP10 induces degradation of oncogenic FLT3. Nature chemical biology 13, 1207–1215, doi:10.1038/nchembio.2486 (2017).

23. Sun, J. et al. USP10 inhibits lung cancer cell growth and invasion by upregulating PTEN. Mol Cell Biochem 441, 1–7, doi:10.1007/s11010-017-3170-2 (2018).

24. Lu, C. et al. USP10 suppresses tumor progression by inhibiting mTOR activation in hepatocellular carcinoma. Cancer letters 436, 139–148, doi:10.1016/j.canlet.2018.07.032 (2018).

25. Zhu, H. et al. USP10 Promotes Proliferation of Hepatocellular Carcinoma by Deubiquitinating and Stabilizing YAP/TAZ. Cancer research 80, 2204–2216, doi:10.1158/0008-5472.CAN-19-2388 (2020).

26. Yuan, J., Luo, K., Zhang, L., Cheville, J. C. & Lou, Z. USP10 regulates p53 localization and stability by deubiquitinating p53. Cell 140, 384–396, doi:10.1016/j.cell.2009.12.032 (2010).

27. Liu, J. et al. Beclin1 controls the levels of p53 by regulating the deubiquitination activity of USP10 and USP13. Cell 147, 223–234, doi:10.1016/j.cell.2011.08.037 (2011).

28. Deng, M. et al. Deubiquitination and Activation of AMPK by USP10. Molecular cell 61, 614–624, doi:10.1016/j.molcel.2016.01.010 (2016).

29. Lim, R. et al. Deubiquitinase USP10 regulates Notch signaling in the endothelium. Science 364, 188-+, doi:10.1126/science.aat0778 (2019).

30. Liao, Y. et al. Inhibition of EGFR signaling with Spautin-1 represents a novel therapeutics for prostate cancer. J Exp Clin Cancer Res 38, 157, doi:10.1186/s13046-019-1165-4 (2019).

31. Koo, B. K. et al. Tumour suppressor RNF43 is a stem-cell E3 ligase that induces endocytosis of Wnt receptors. Nature 488, 665–669, doi:10.1038/nature11308 (2012).

32. Chen, M. et al. TMEM79/MATTRIN defines a pathway for Frizzled regulation and is required for Xenopus embryogenesis. eLife 9, doi:10.7554/eLife.56793 (2020).

33. Madan, B. et al. USP6 oncogene promotes Wnt signaling by deubiquitylating Frizzleds. Proceedings of the National Academy of Sciences of the United States of America 113, E2945–2954, doi:10.1073/pnas.1605691113 (2016).

34. Lu, Y. et al. Twa1/Gid8 is a beta-catenin nuclear retention factor in Wnt signaling and colorectal tumorigenesis. Cell research 27, 1422–1440, doi:10.1038/cr.2017.107 (2017).

35. Novellasdemunt, L. et al. USP7 Is a Tumor-Specific WNT Activator for APC-Mutated Colorectal Cancer by Mediating beta-Catenin Deubiquitination. Cell reports 21, 612–627, doi:10.1016/j.celrep.2017.09.072 (2017).

36. Tanneberger, K. et al. Structural and functional characterization of the Wnt inhibitor APC membrane recruitment 1 (Amer1). The Journal of biological chemistry 286, 19204–19214, doi:10.1074/jbc.M111.224881 (2011).

37. Rivera, M. N. et al. An X chromosome gene, WTX, is commonly inactivated in Wilms tumor. Science 315, 642–645, doi:10.1126/science.1137509 (2007).

38. Major, M. B. et al. Wilms tumor suppressor WTX negatively regulates WNT/beta-catenin signaling. Science 316, 1043–1046, doi:10.1126/science/1141515 (2007).

39. Schwarz-Romond, T. et al. The DIX domain of Dishevelled confers Wnt signaling by dynamic polymerization. Nature structural & molecular biology 14, 484–492, doi:10.1038/nsmb1247 (2007).

40. Musacchio, A. On the role of phase separation in the biogenesis of membraneless compartments. EMBO J 41, e109952, doi:10.15252/embj.2021109952 (2022).

41. Tiwary, A. K. & Zheng, Y. X. Protein phase separation in mitosis. Current Opinion in Cell Biology 60, 92–98, doi:10.1016/j.ceb.2019.04.011 (2019).

42. Davis, R. B., Moosa, M. M. & Banerjee, P. R. Ectopic biomolecular phase transitions: fusion proteins in cancer pathologies. Trends Cell Biol 32, 681–695, doi:10.1016/j.tcb.2022.03.005 (2022).

43. Banani, S. F., Lee, H. O., Hyman, A. A. & Rosen, M. K. Biomolecular condensates: organizers of cellular biochemistry. Nat Rev Mol Cell Bio 18, 285–298, doi:10.1038/nrm.2017.7 (2017).

44. Li, Y. et al. Volumetric Compression Induces Intracellular Crowding to Control Intestinal Organoid Growth via Wnt/beta-Catenin Signaling. Cell Stem Cell 28, 63–78 e67, doi:10.1016/j.stem.2020.09.012 (2021).

45. Martino-Echarri, E., Brocardo, M. G., Mills, K. M. & Henderson, B. R. Tankyrase Inhibitors Stimulate the Ability of Tankyrases to Bind Axin and Drive Assembly of beta-Catenin Degradation-Competent Axin Puncta. PloS one 11, e0150484, doi:10.1371/journal.pone.0150484 (2016).

46. Schneider, S., Steinbeisser, H., Warga, R. M. & Hausen, P. beta-Catenin translocation into nuclei demarcates the dorsalizing centers in frog and fish embryos. Mech Develop 57, 191–198, doi:Doi 10.1016/0925-4773(96)00546-1 (1996).

47. Parker, D. S., Ni, Y. Y., Chang, J. L., Li, J. & Cadigan, K. M. Wingless signaling induces widespread chromatin remodeling of target loci. Molecular and cellular biology 28, 1815–1828, doi:10.1128/MCB.01230-07 (2008).

48. Kelly, C., Chin, A. J., Leatherman, J. L., Kozlowski, D. J. & Weinberg, E. S. Maternally controlled beta-catenin-mediated signaling is required for organizer formation in the zebrafish. Development 127, 3899–3911 (2000).

49. Dorsky, R. I., Sheldahl, L. C. & Moon, R. T. A transgenic Lef1/beta-catenin-dependent reporter is expressed in spatially restricted domains throughout zebrafish development. Dev Biol 241, 229–237, doi:10.1006/dbio.2001.0515 (2002).

50. Sato, T. et al. Single Lgr5 stem cells build crypt-villus structures in vitro without a mesenchymal niche. Nature 459, 262–265, doi:10.1038/nature07935 (2009).

51. Almeqdadi, M., Mana, M. D., Roper, J. & Yilmaz, O. H. Gut organoids: mini-tissues in culture to study intestinal physiology and disease. Am J Physiol Cell Physiol 317, C405–C419, doi:10.1152/ajpcell.00300.2017 (2019).

52. Sato, T. & Clevers, H. Growing self-organizing mini-guts from a single intestinal stem cell: mechanism and applications. Science 340, 1190–1194, doi:10.1126/science.1234852 (2013).

53. Zhan, T., Rindtorff, N. & Boutros, M. Wnt signaling in cancer. Oncogene 36, 1461–1473, doi:10.1038/onc.2016.304 (2017).

54. He, Y. et al. The deubiquitinase USP10 restores PTEN activity and inhibits non-small cell lung cancer cell proliferation. The Journal of biological chemistry 297, 101088, doi:10.1016/j.jbc.2021.101088 (2021).

55. Kim, K. et al. Prognostic significance of USP10 and p14ARF expression in patients with colorectal cancer. Pathol Res Pract 216, 152988, doi:10.1016/j.prp.2020.152988 (2020).

56. Wang, X. et al. The deubiquitinase USP10 regulates KLF4 stability and suppresses lung tumorigenesis. Cell Death Differ 27, 1747–1764, doi:10.1038/s41418-019-0458-7 (2020).

57. Dennler, S. et al. Direct binding of Smad3 and Smad4 to critical TGF beta-inducible elements in the promoter of human plasminogen activator inhibitor-type 1 gene. EMBO J 17, 3091–3100, doi:10.1093/emboj/17.11.3091 (1998).

58. Schneider, C. A., Rasband, W. S. & Eliceiri, K. W. NIH Image to ImageJ: 25 years of image analysis. Nat Methods 9, 671–675, doi:10.1038/nmeth.2089 (2012).

59. Kimmel, C. B., Ballard, W. W., Kimmel, S. R., Ullmann, B. & Schilling, T. F. Stages of embryonic development of the zebrafish. Developmental dynamics : an official publication of the American Association of Anatomists 203, 253–310, doi:10.1002/aja.1002030302 (1995).

60. Wei, S. et al. The guanine nucleotide exchange factor Net1 facilitates the specification of dorsal cell fates in zebrafish embryos by promoting maternal beta-catenin activation. Cell Res 27, 202–225, doi:10.1038/cr.2016.141 (2017).

61. Koo, B. K., Sasselli, V. & Clevers, H. Retroviral gene expression control in primary organoid cultures. Curr Protoc Stem Cell Biol 27, Unit 5A 6, doi:10.1002/9780470151808.sc05a06s27 (2013).

62. Cai, Z. L. et al. Optimized digital counting colonies of clonogenic assays using ImageJ software and customized macros: Comparison with manual counting. Int J Radiat Biol 87, 1135–1146, doi:10.3109/09553002.2011.622033 (2011).

