## Supplementary material for "USP10 strikes down β-catenin by dual-wielding deubiquitinase activity and phase separation potential": None

### Supplemental Information

#### Supplemental figure

#### Supplemental figure legends

### Supplementary Figure 1

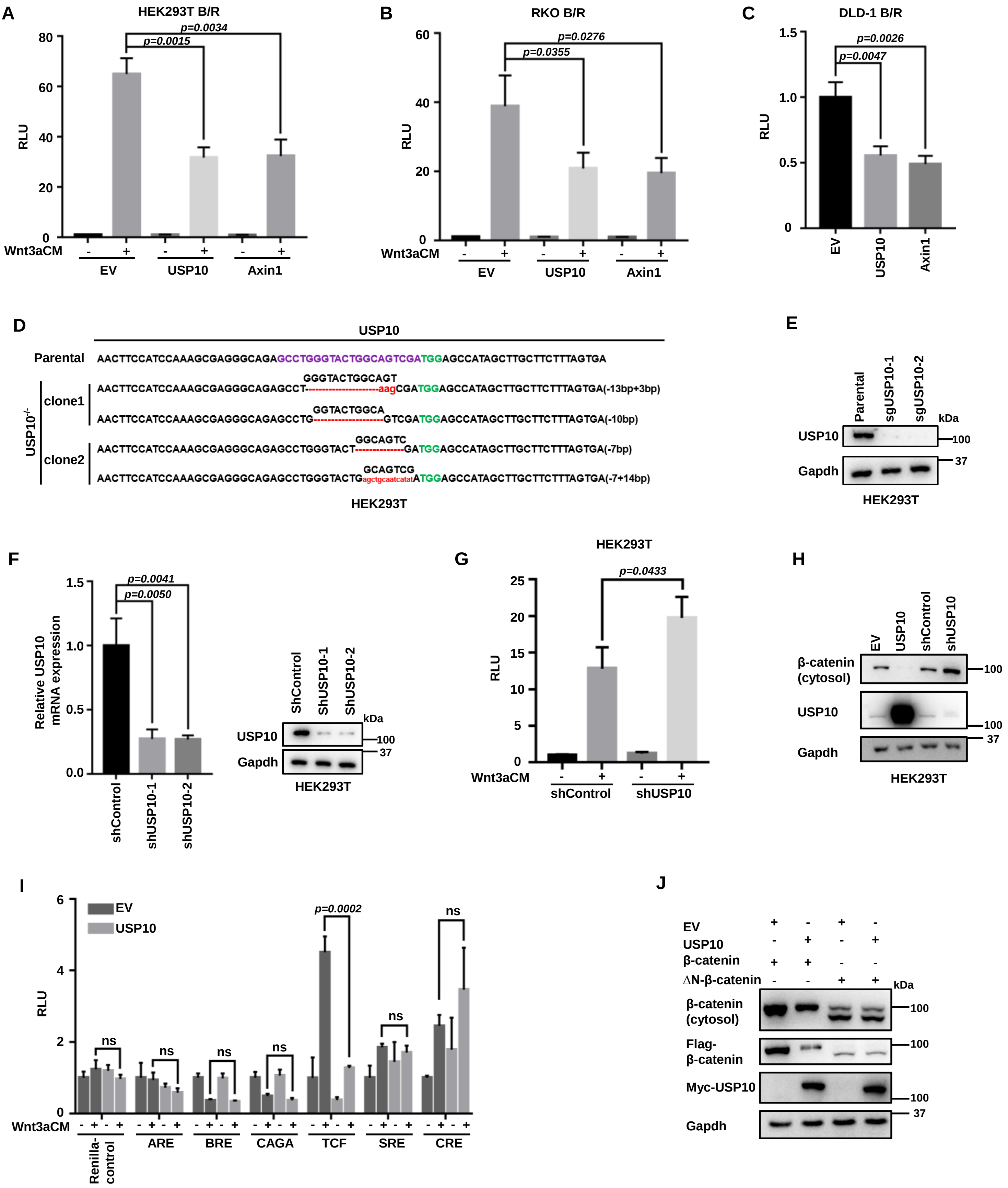

**Supplemental Figure 1.** Additional experimental results demonstrate USP10 inhibits Wnt/β-catenin signaling, related to **Figure 1**.

(A) TOPFlash reporter assay showing USP10 inhibits Wnt/β-catenin signaling in HEK293T cell. Axin1 is used as positive control. Error bars mean ± SD, n = 3, two-tailed Student's t-test.

(B) TOPFlash reporter assay showing USP10 inhibits Wnt/β-catenin signaling in RKO cell. Axin1 is used as positive control. Error bars mean ± SD, n = 3, two-tailed Student's t-test.

(C) TOPFlash reporter assay showing USP10 inhibits Wnt/β-catenin signaling in DLD-1 cell in the absence of exogenous Wnt. Axin1 is used as positive control. Error bars mean ± SD, n = 3, two-tailed Student's t-test.

(D) Sanger sequence of the two clones of USP10 CRISPR-Cas9 knockout in HEK293T cells.

(E) Validation of USP10 knockout by WB.

(F) Validation of USP10 shRNA knockdown by RT-qPCR and WB. Error bars mean ± SD, n = 3, two-tailed Student's t-test. shUSP10-1 was used and abbreviated as shUSP10 in sequel experiments.

(G) Knockdown of USP10 significantly enhances TOPFlash reporter at the presence of Wnt3a CM. Error bars mean ± SD, n = 3, two-tailed Student's t-test.

(H) WB assay showing the alteration of cytosolic β-catenin levels under USP10 overexpression or knockdown condition.

(I) Dual-luciferase chemiluminescence of ARE, BRE, CAGA, TCF, SRE and CRE under control CM or Wnt3a CM treatment. Error bars mean ± SD, n = 3, two-tailed Student's t-test.

(J) WB assay showing USP10 overexpression cannot effectively reduce the cytosolic ΔN-β-catenin level.

RLU: Relative Luciferase Unit. ns, not significant.

### Supplementary Figure 2

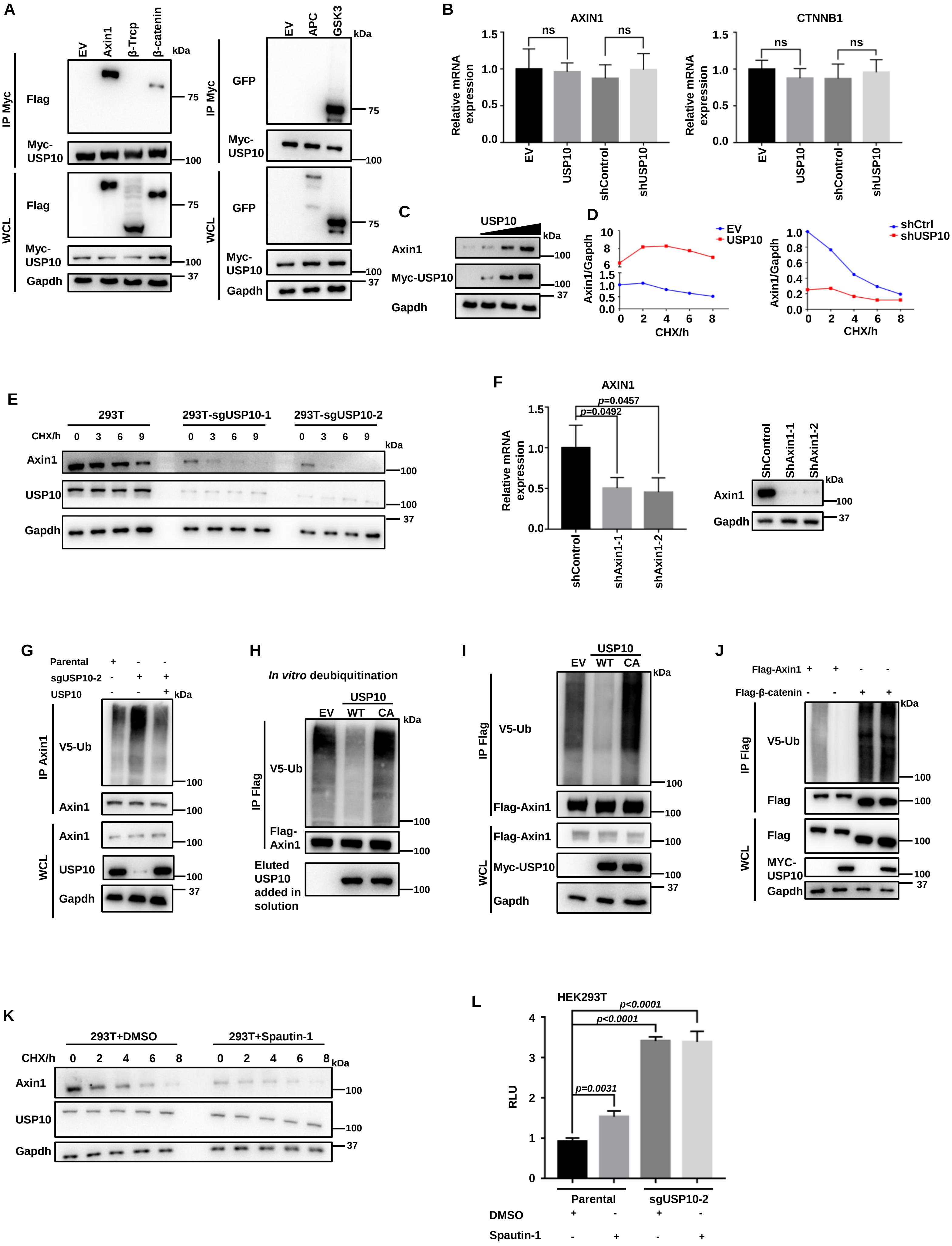

**Supplemental Figure 2.** Supportive information for deubiquitination and stabilization of Axin1 by USP10, related to **Figure 2**.

(A) Co-IP of USP10 with exogenous key components of β-catenin destruction complex, including Axin1, β-Trcp, β-catenin, GSK3 and APC.

(B) Validation of USP10 knockdown effect on AXIN1 and CTNNB1 transcription levels. Error bars mean ± SD, n = 3, two-tailed Student's t-test.

(C) WB showing endogenous Axin1 level increases with USP10 dose-dependently.

(D) Quantifications of Axin1 levels in the pulse-chase assay shown by Figure 2c.

(E) Pulse-chase assay of endogenous Axin1 under USP10 KO conditions.

(F) Validation of human Axin1 knockdown by RT-qPCR and WB. shAxin1-1 was used and abbreviated as shAxin1 in sequel experiments. Error bars mean ± SD, n = 3, two-tailed Student's t-test.

(G) Ubiquitination assay of Axin1 under USP10 knockout and rescue conditions.

(H) *In vitro* deubiquitination assay of USP10 on Axin1.

(I) Ubiquitination assay showing USP10-CA mutant loses the capability to deubiquitinate exogenous Axin1.

(J) Ubiquitination assay showing USP10 does not deubiquitinate β-catenin.

(K) Pulse-chase assay of endogenous Axin1 under Spautin-1 treatment.

(L) TOPFlash reporter assay in parental or USP10 KO HEK293T cell under Spautin-1 treatment. Error bars mean ± SD, n = 3, two-tailed Student's t-test.

### Supplementary Figure 3

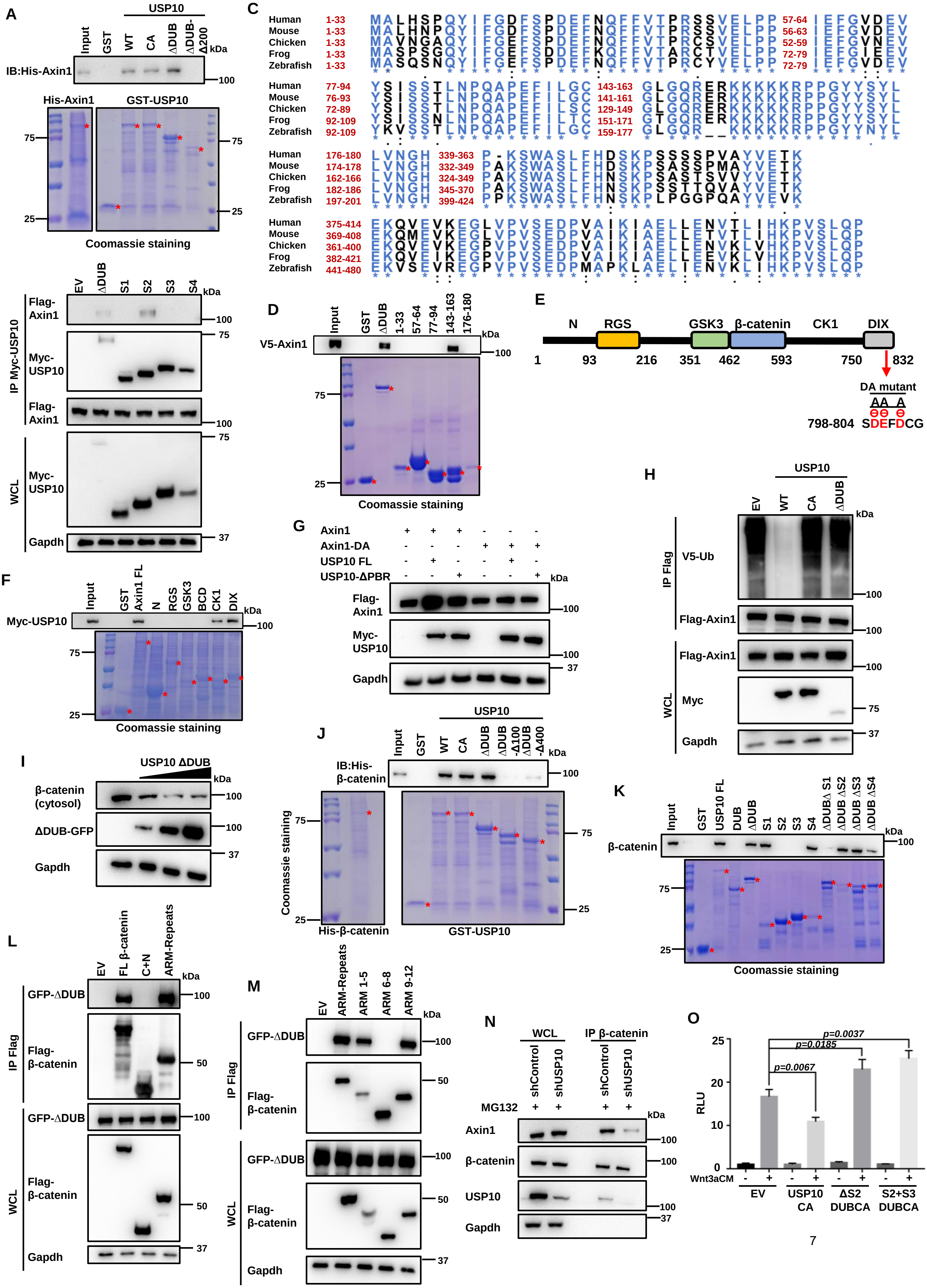

**Supplemental Figure 3.** Supportive information for Axin1 and β-catenin binding to USP10, related to **Figure 3**.

(A) *In vitro* pull-down assay of bacterial expressed Axin1 and USP10. Asterisks represent the major bands of the desired proteins.

(B) Co-IP assay of different segments of USP10 with Axin1.

(C) Amino acid sequence alignment showing the seven conserved regions on ΔDUB of vertebrate USP10.

(D) Pull-down assay of Axin1 with different conserved regions of USP10-ΔDUB. Asterisks represent the major bands of the desired proteins.

(E) Schematic drawing of Axin1 protein. N, N-terminal; RGS, RGS domain; GSK3, GSK3 binding helix; β-cat, β-catenin binding site; CK1, CK1 putative binding site; DIX, DIX domain.

(F) Pull-down assay of USP10 with different regions of Axin1. Asterisks represent the major bands of the desired proteins.

(G) Axin1 WT/DA levels alterations under USP10 WT or ΔPBR overexpression condition.

(H) Ubiquitination assay of Axin1 under USP10 WT, CA, and ΔDUB overexpression.

(I) WB assay showing cytosolic β-catenin level diminishes dose-dependently of USP10-ΔDUB.

(J) *In vitro* pull-down assay of bacterial expressed β-catenin and USP10. Asterisks represent the major bands of the desired proteins.

(K) Pull-down assay showing different segments of USP10 interacting with β-catenin.

(L, M) Co-IP assay of USP10 ΔDUB with different segments of β-catenin.

(N) Co-IP assay showing depletion of USP10 results in the reduced interaction between endogenous Axin1 and β-catenin.

(O) TOPFlash reporter assay showing USP10 ΔS2+DUBCA and USP10 S2+S3+DUBCA behave dominant-negatively. Error bars mean ± SD, n = 3, two-tailed Student's t-test.

RLU: Relative Luciferase Unit. ns, not significant.

### Supplementary Figure 4

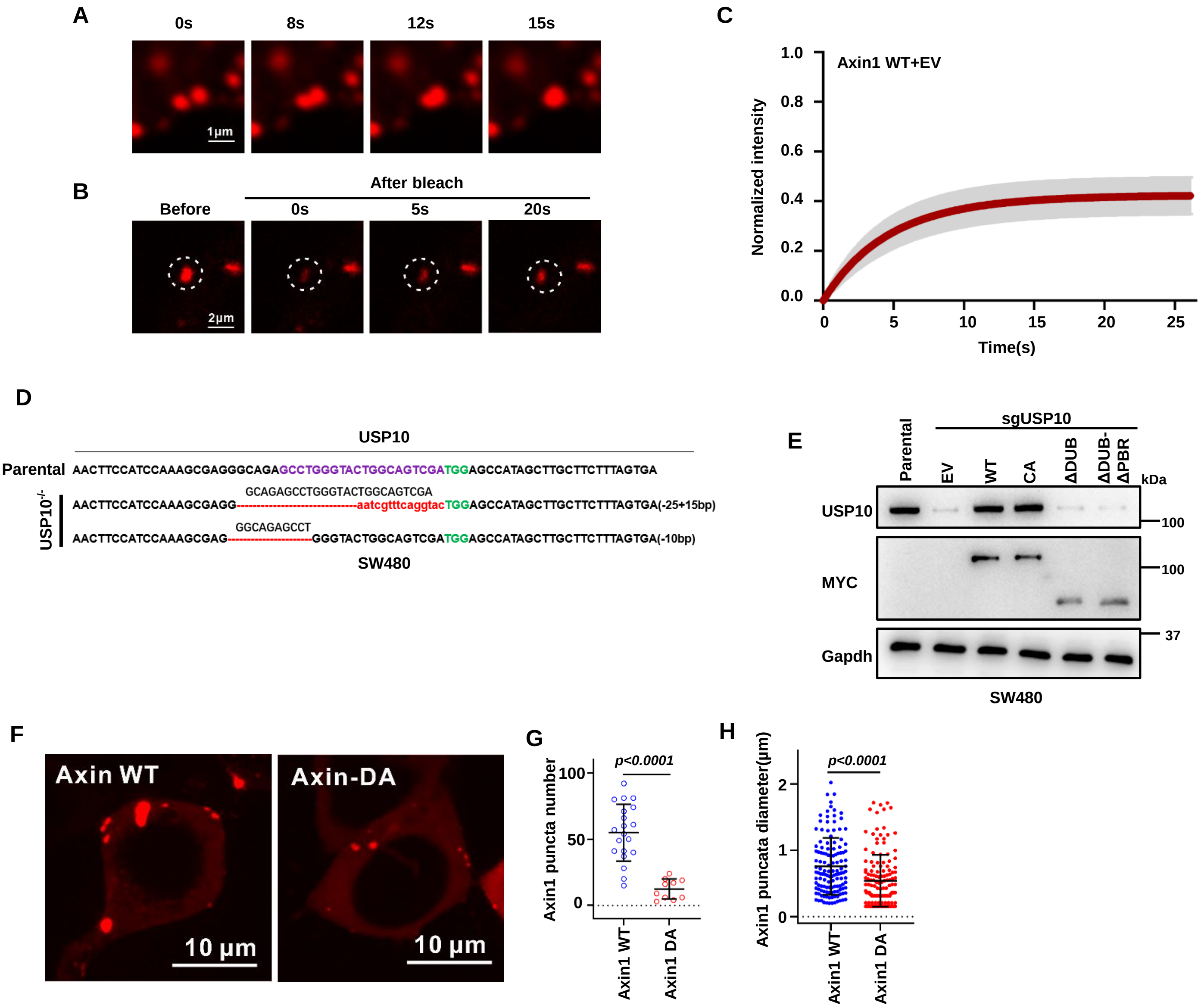

**Supplemental Figure 4.** Supportive information showing USP10 promotes Axin1 granule formation, related to **Figure 4**.

(A) Representative fluorescent images showing the fusion of Axin1 puncta.

(B) Representative images of the FRAP assay of Axin1 puncta.

(C) Recovery of fluorescent intensity after the photobleaching during the FRAP assay of Axin1 puncta.

(D) Sanger sequence of the USP10 CRISPR-Cas9 knockout in SW480 cells.

(E) Protein level validation of USP10 KO and rescue in SW480 in Figure 4h.

(F) Representative fluorescent images of Axin1 droplets (WT and DA).

(G, H) Numbers of Axin1 puncta per cell (G), size of Axin1 puncta (H). Error bars mean ± SD, two-tailed Student's t-test.

### Supplementary Figure 5

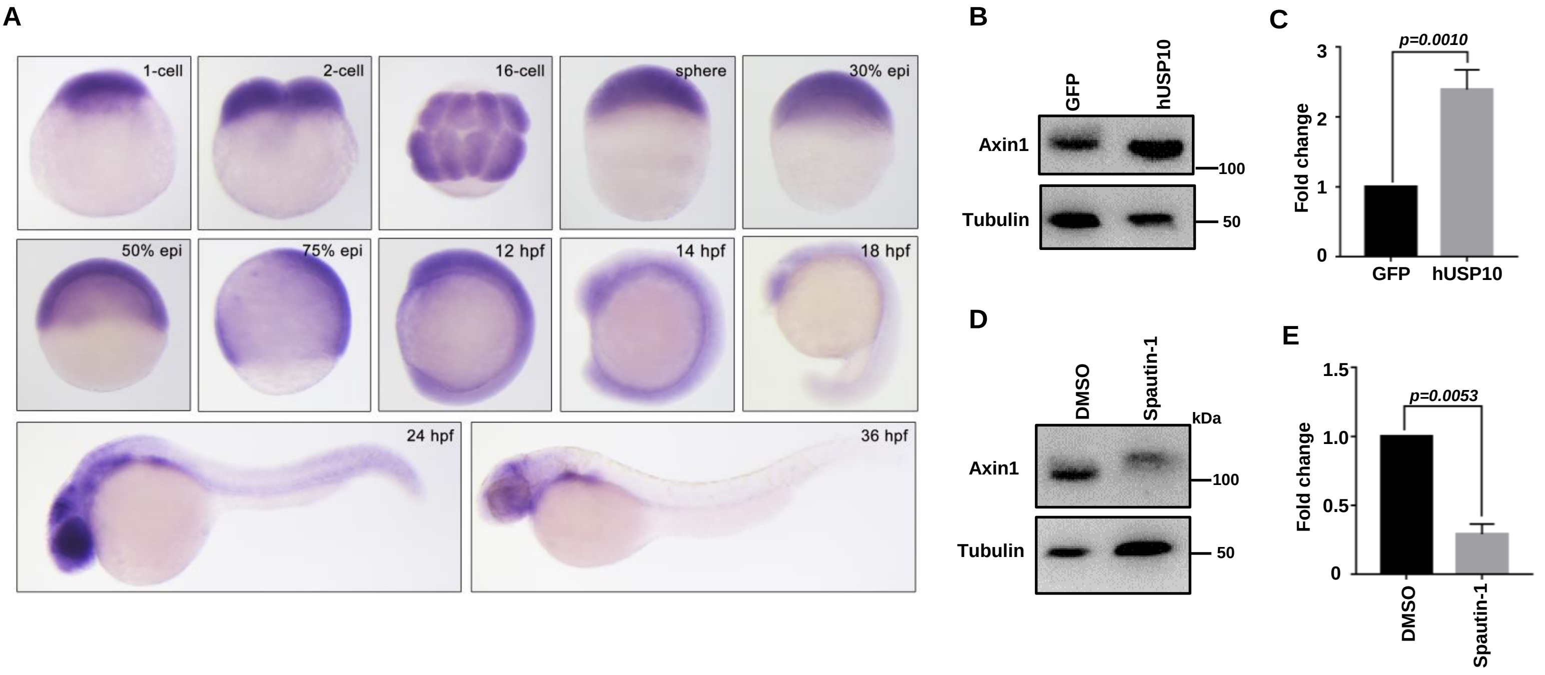

**Supplementary Figure 5.** Supportive assays of zebrafish embryo study, related to **Figure 5**.

(A) Expression analysis of the zebrafish *usp10* transcript at the indicated stages by whole-mount *in situ* hybridization. Embryos were shown with lateral view except the 16-cell stage (animal pole views).

(B-E) USP10 promoted the stabilization of Axin1. The hUSP10 overexpression and Spautin-1 treated embryos were collected at sphere stage for WB with the indicated antibodies (B and D) and the protein levels were analyzed respectively in (C and E). Error bars mean ± SD, n = 3, two-tailed Student's t-test.

#### Supplementary Figure 6

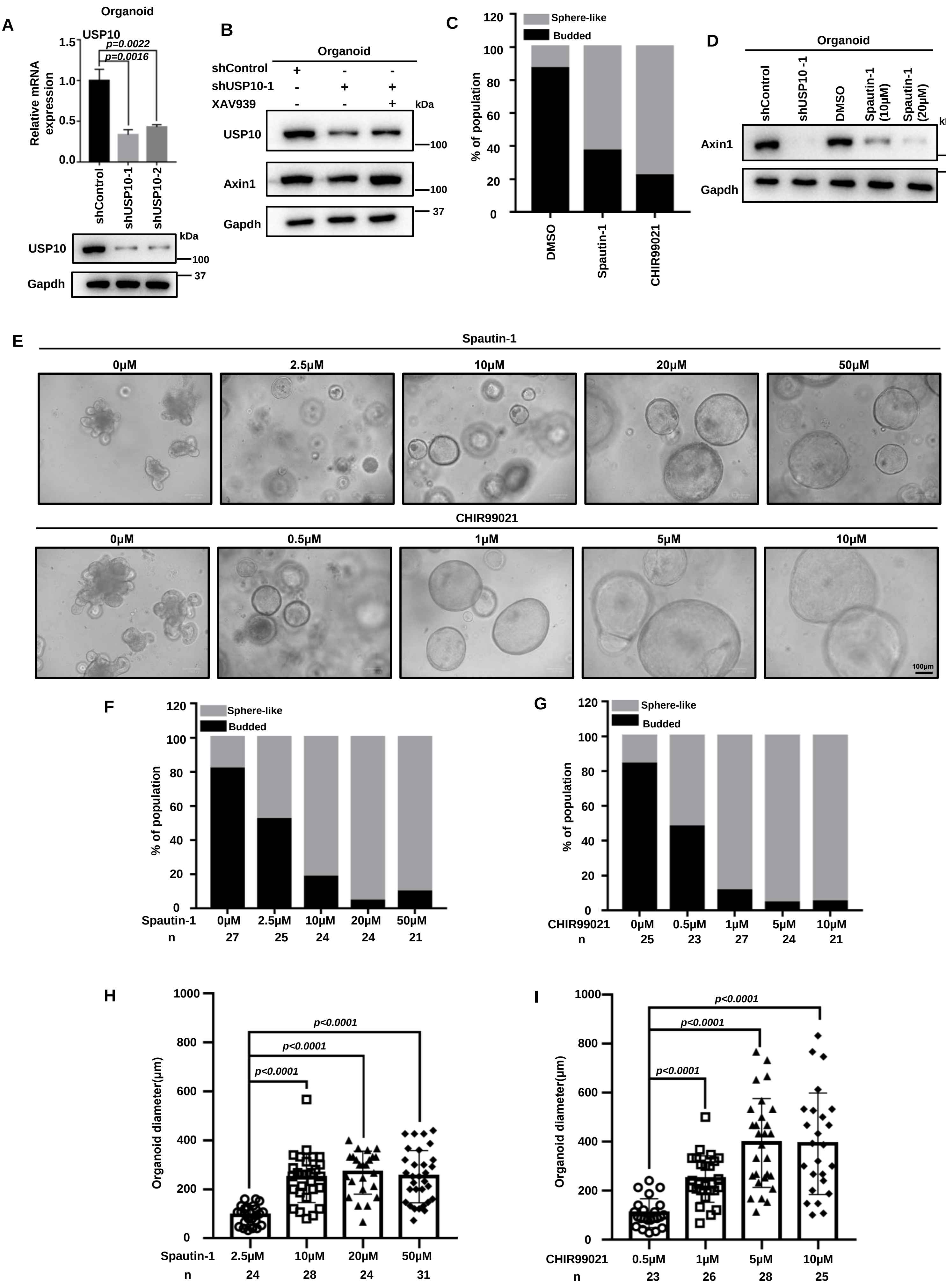

**Supplemental Figure 6.** Supportive assays of intestinal organoid study, related to **Figure 6**.

(A) Validation of murine USP10 knockdown by RT-qPCR and WB in the organoids. shUSP10-1 was used and abbreviated as shUSP10 in the sequel experiments.

(B) WB showing USP10 and Axin1 levels under USP10 depletion and/or XAV939 treatment.

(C) Percentage of sphere-like and budded organoids under Spautin-1 and CHIR99021 treatments. DMSO group:n=45, Spautin-1 group:n=46, CHIR99021 group:n=49.

(D) WB assay showing treatment of Spautin-1 effectively reduced endogenous Axin1 level in murine intestinal organoids. shUSP10 was used as control to confirm the effect of Spautin-1.

(E) Representative images of murine intestinal organoids under DMSO, Spautin-1 or CHIR99021 treatment at different doses. All images in the panel are in the same scale.

(F, G) Percentage of sphere-like and budded organoids under Spautin-1 and CHIR99021 treatments.

(H, I) Organoid (sphere-like) diameters quantifications in (E) shown by histogram. Error bars mean ± SD, by two-tailed Student's t-test.

### Supplementary Figure 7

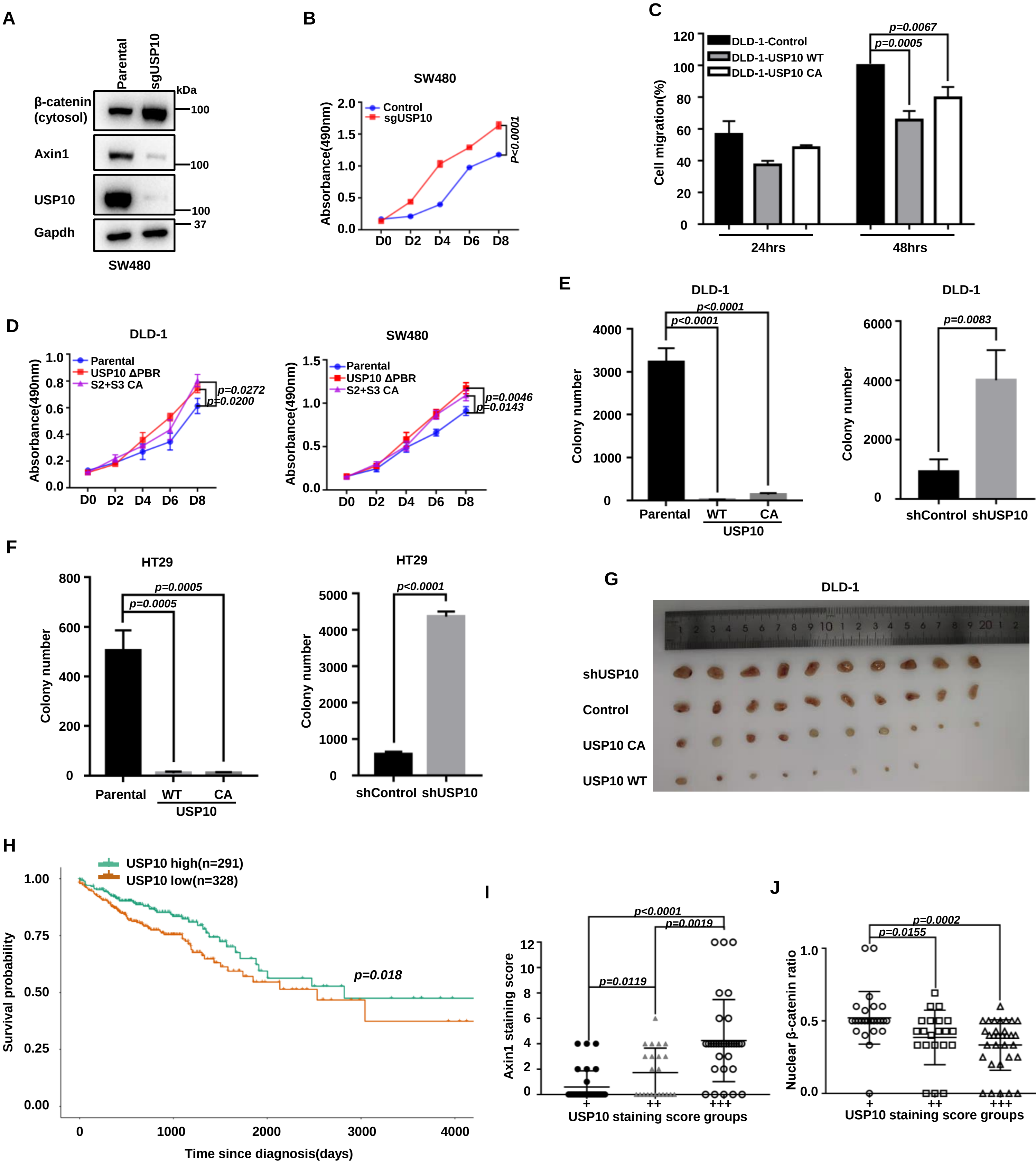

**Supplemental Figure 7.** Additional assays and information to illustrate the tumor-suppressive effect of USP10 in human CRC, related to **Figure 7**.

(A) Endogenous Axin1 and cytosolic β-catenin levels in SW480 cell lines under USP10 Knockout condition.

(B) MTT assay showing USP10 KO significantly enhanced SW480 cell growth. Error bars mean ± SD, n = 3, two-way ANOVA.

(C) Wound healing assay of DLD-1 cell after expression of USP10 WT and USP10-CA. Error bars mean ± SD, n = 3, two-tailed Student's t-test.

(D) MTT cell growth assay of DLD-1 and SW480 cells with the overexpression of dominant negative USP10 mutants. Error bars mean ± SD, n = 3, two-way ANOVA.

(E, F) Quantifications of the numbers of colonies in Figure 7F-G. Error bars mean ± SD, by two-tailed Student's t-test.

(G) Images of the tumors formed by subcutaneously transplanted DLD-1 cells after abscission.

(H) Overall survival of USP10-high and -low patients from TCGA-COADREAD database.

(I, J) Correlation analysis between Axin1 and nuclear localized β-catenin ratio based on grouped USP10 levels. Error bars mean ± SD, n = 3, two-tailed Student's t-test. ns, not significant.
